# Latitudinal diversity gradients and selective microbial exchange at the Atlantic ocean–air interface

**DOI:** 10.64898/2026.03.24.713978

**Authors:** Isabella Hrabe de Angelis, S. Emil Ruff, Hedy M. Aardema, Sanja Basic, Jan D. Brüwer, Hans A. Slagter, Jens Weber, Maria Ll. Calleja, Zoe G. Cardon, Antonis Dragoneas, Anna Lena Leifke, Björn Nillius, Subha S. Raj, David Walter, Bettina Weber, Bernhard M. Fuchs, Gerald H. Haug, Ulrich Pöschl, Ralf Schiebel, Christopher Pöhlker

**Affiliations:** Max Planck Institute for Chemistry, Multiphase Chemistry Department, Hahn-Meitner-Weg 1, 55128 Mainz, Germany; Marine Biological Laboratory, Bay Paul Center, Woods Hole, MA 02543, USA; Marine Biological Laboratory, Ecosystems Center, Woods Hole, MA 02543, USA; Max Planck Institute for Chemistry, Climate Geochemistry Department, Hahn-Meitner-Weg 1, 55128 Mainz, Germany; ETH Zurich, Department of Earth Sciences, Sonneggstrasse 5, 8092 Zurich, Switzerland; Max Planck Institute for Marine Microbiology, Celsiusstraße 1, 28359 Bremen, Germany; University of Graz, Institute of Biology, Holteigasse 6, 8010 Graz, Austria; University of the Balearic Islands, Marine Ecology and Systematics Group, Cra. Valldemossa Km. 7.5, 07122 Palma de Mallorca, Spain

## Abstract

Microbial communities at the ocean-atmosphere interface play a vital role in nutrient, aerosol, and water cycling, yet their large-scale biogeography remains scarcely studied. Here we assessed microbial cell abundance and community structure in air and surface ocean samples along a 14,400 km transect from the polar circle to the equator. Air and surface ocean microbiomes were taxonomically distinct across the North East Atlantic. The microbiome in the air had a lower local community richness compared to the surface ocean, yet was more diverse across larger geographical scales. Our results suggest that terrestrial-derived air masses affected air and surface ocean communities. Air microbial communities showed a latitudinal diversity gradient with richness and cell abundance increasing from the polar circle to the equator. We observed an exchange of microbial lineages between ocean and air, however, most lineages were confined to one realm, suggesting lineage-selective aerosolization and deposition of microbial cells.

## Introduction

Earth’s atmosphere represents a vast environment for microorganisms (1), and aerosol of biological origin can represent up to 25 % of the total aerosol mass (2). Average atmospheric cell abundances are considered to be lower over the ocean (∼1 × 10^4^ cells m^−3^) compared to land (1.2 × 10^4^ to 6.5 × 10^5^ cells m^−3^). Yet, the relative contribution of microorganisms to the total aerosol load, and hence their relative importance for atmospheric processes, increases with decreasing particle concentrations at remote locations over the open ocean (3). The world’s oceans, covering more than two thirds of the Earth’s surface, are thus a major, yet poorly understood global aerosol source (4). Depending on sampling location and conditions, microbial cell concentrations over the oceans range from ∼10^1^ to 10^6^ cells m^−3^ (3; 5; 6; 7; 8; 9). Through large-scale initiatives, such as the Tara Oceans project, the Antarctic Circumnavigation Expedition, and the Malaspina 2010 circumnavigation, our understanding of marine aerosols on a global scale has improved, showing distinct microbiomes in ocean and air (6; 10; 11). The marine air microbiome is highly variable, driven by the interplay of marine, terrestrial, natural, and anthropogenic sources (12). A solid understanding of the formation and exchange of bioaerosols is crucial to assess their impact on the climate system and global biogeography and has become a topic of increasing interest in marine ecology and climate studies (10; 11; 13; 14; 15; 16).

The extent to which particles and microorganisms are aerosolized from the ocean, as well as their residence time in the air, depend on several factors, including surface wind speed (17), particle size and chemistry (18) and sea state, i.e., wave direction, height and shape (19; 20). Once aerosolized, particles can stay airborne up to several days and can be transported over long distances (6). Microbes found in air at remote locations are often associated with dust particles (21; 22; 23). These airborne particles can affect the hydrological cycle (cloud formation and precipitation) by acting as cloud condensation nuclei or ice nuclei. Microbes can act as ice nucleation particles, or express proteins triggering ice nucleation (24), already being effective at temperatures close to 0 °C (25).

Particles that are dry- or wet-deposited into a new ecosystem, may introduce nutrients and biodiversity to habitats, far away from the source location (13; 26; 27). The introduction of nutrients, pollutants, trace metals and microbial communities from terrestrial sources can be especially relevant in the nutrient-limited oligotrophic oceans (28). Marine environments were highlighted as regions of key interest for bioaerosol research (29). Understanding the effect of the atmospheric transport of nutrients and terrestrial microbial communities on the surface ocean microbiome is critical to assess ocean productivity and element cycling, and to estimate potential impacts of changes in land-use, aridification and extreme weather events (30).

Insights into open ocean air microbiomes are often hampered by a lack of continuous or large-scale data, by particle contamination from ship exhaust, by biological contamination during sampling, and by extremely low biomass samples (31). With the research sailing vessel *Eugen Seibold* (32), we had the unique opportunity to sample air across large geographical scales, using rigorous biological contamination controls and unaffected by the ship’s exhaust. With the vessel’s specialized instrumentation to focus on sea-air interface processes we studied the microbiology and chemistry of the atmospheric boundary layer at unprecedented resolution. We investigated the origin, abundance, composition, dispersal, and exchange of microbial communities in the lower atmosphere and surface water of the North East Atlantic Ocean. We sampled six marine geographical provinces (33) along a latitudinal transect from 67° N (polar circle) to 3° N (equator), roughly following the 20° W meridian. Meteorological, oceanographic, biochemical and physical measurements, as well as satellite observations complement the data set. We highlight fundamental differences between seawater and air microbiomes, explore the impact of long-range transport from continents, compare terrestrial-influenced with pristine marine conditions, and assess the importance and selectivity of microbial air-sea exchange.

## Results and Discussion

### Air and surface ocean microbial diversity significantly differ in the North East Atlantic Ocean

Along a 14,400 km long latitudinal transect in the North East Atlantic, we analyzed the microbial communities of the marine atmospheric boundary layer (∼3.5 m above the ocean surface), and the surface ocean water (∼3 m water depth; Fig. 1, Fig. S1). We collected and analyzed a total of 77 samples, 38 air samples and 39 water samples from 40 different sites, with paired ocean-atmosphere sampling at 36 sites (Dataset S1). We sequenced archaeal and bacterial 16S rRNA gene amplicons for each sample and collected a comprehensive set of atmospheric and oceanic physicochemical and biological parameters at each site.

**Figure 1.**
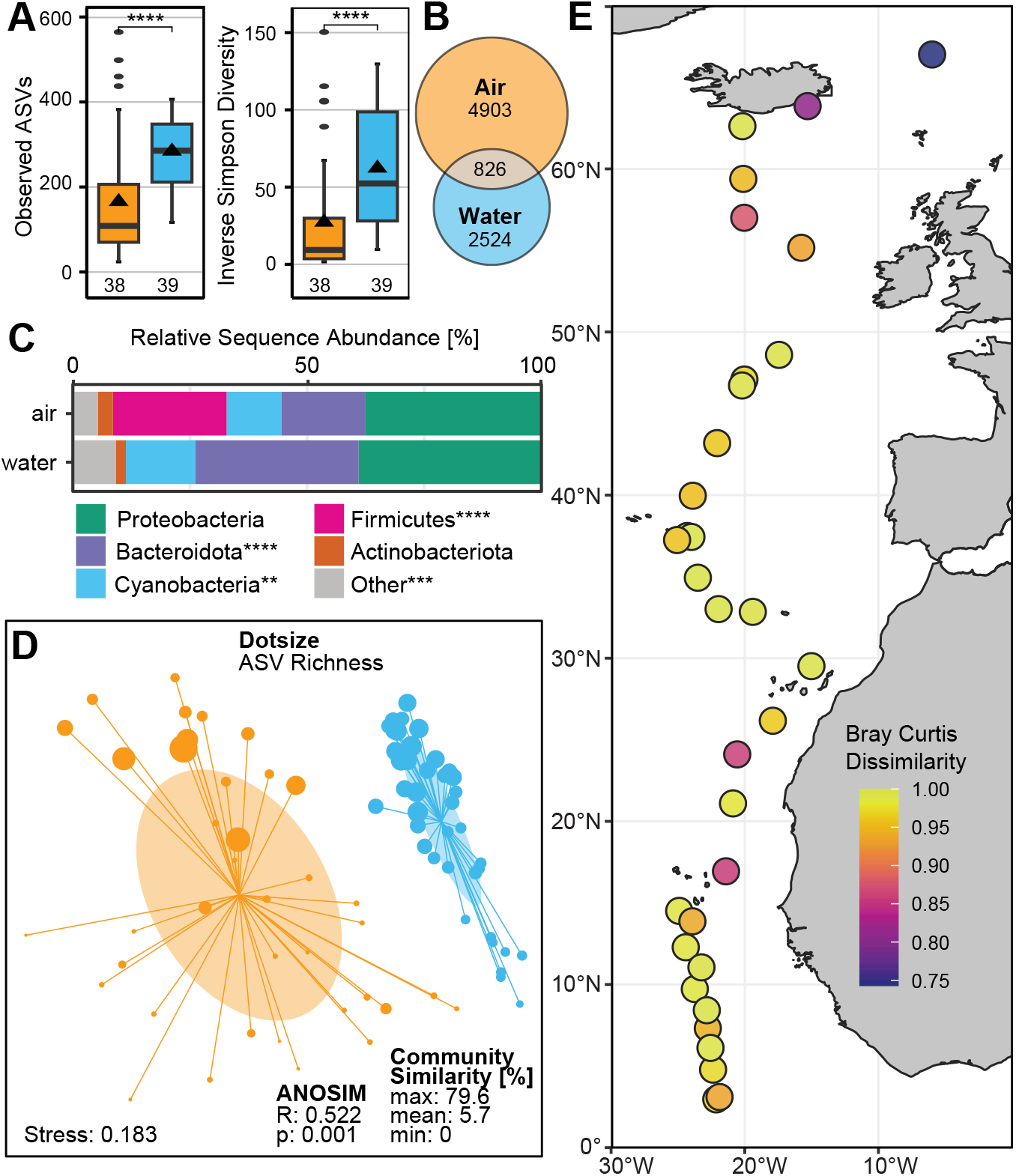
Microbial diversity of lower atmosphere and surface ocean communities of the North East Atlantic Ocean. A: Microbial alpha diversity of air (orange, n_Samples_ = 38) and water (blue, n_Samples_ = 39) samples. Boxplots show mean (triangle), median (middle line), upper and lower quartile (top and lower line). Whiskers show maximum data points within upper/lower quartile plus 1.5 times the interquartile range (IQR). Dots represent outlier values that exceed 1.5 × IQR. Groups were tested using a Wilcoxon rank sum test, ****: p<0.0001. B: Scaled Venn Diagram showing unique and shared ASVs between surface ocean and air, corrected for group size. C: Relative sequence abundances of the five most abundant phyla. Differences between air and water were tested using a Wilcoxon rank sum test, ****: p<0.0001, ***: p<0.001, **: p<0.01. D: Non-metric multidimensional scaling (NMDS) plot of air (orange) and surface ocean (blue) communities. The closer two dots are, the more similar are their underlying communities. Dot size scales with sample richness, oval shapes depict one standard deviation of the centroid (weighted average within-group-distance). Lines connect each dot to the centroid of the tested group. E: Bray Curtis dissimilarities between microbial communities of water vs air sample pairs along the transect (0: identical communities, 1: completely different communities).

The amplicon sequence variant (ASV) based diversity and composition of microbial communities differed significantly between air and surface ocean along the entire transect (Fig. 1A-E). Alpha diversity, i.e., local or per sample diversity, including richness (number of unique ASVs) and evenness (proportion of ASVs), was significantly lower in air communities compared to ocean communities (Fig. 1A, Wilcoxon rank sum test: *p*<0.0001, Dataset S1).

Within the air and ocean biomes, air samples shared fewer ASVs among each other, i.e., had a higher community turnover (beta diversity, differences between locations), than water samples (Fig. 1D, Fig. S2). Thus, air communities were lower in richness but differed strongly between locations, while ocean communities had a higher richness, with low spatial variability. Greater community differences in the air, compared to the water, seem to be a conserved pattern for the North Atlantic and Western Pacific Ocean (10). A lower alpha diversity in air samples, however, has so far only been described for the Southern Ocean surrounding Antarctica (11).

The observed large community differences between air masses may be caused by diversity loss due to localized wet deposition, limited mixing between sites, or mixing may be obscured by the heterogeneity of aerosols deriving from various marine and terrestrial sources (34). The observed similarity of surface ocean communities is likely a result of regional mixing and transport due to oceanic currents (35).

Gamma diversity (total diversity found in all samples of a biome) was higher in the air than in the water (Fig. 1B). Overall, 90 % of all ASVs were unique to either the air or the ocean microbiome (normalized to group size). The stark difference between air and ocean communities is also evident using distance-based methods and is corroborated by pairwise Bray-Curtis dissimilarities of >0.74 between the community structures of air and water sample pairs (Fig. 1E, Dataset S1).

The considerable differences between air and ocean community structure (Fig. 1A&D), together with the high proportion of unique ASVs in each biome (Fig. 1B) show that air and water masses harbor fundamentally different habitats and niches. The differences between air and water concerning, e.g., viscosity, water activity, nutrients, organic carbon and radiation exposure seem to select for distinct communities in each realm.

### Taxonomic diversity reveals indicator clades for air and surface ocean communities

Although airborne microbes are typically considered as transient and in exchange with surfaces (1), indicator clades could offer evidence to assess whether air masses serve as a microbial habitat (31), or support the finding that certain clades are preferentially aerosolized or better adapted to airborne conditions, and thus constituting a larger portion of a transient air microbiome.

The North East Atlantic Ocean air samples contained 40, and the oceanic samples 23 phyla. Of these, 21 were unique for the air, 4 were unique for the ocean community, and 19 were shared (Fig. S3&5, Dataset S2&3). The air microbiome was dominated by Proteobacteria (Pseudomonadota, 38 % ± 28 %, average relative sequence abundance (RSA) ± standard deviation), Firmicutes (Bacillota, 24 % ± 31 %), Bacteroidota (18 % ± 19 %), Cyanobacteria (12 % ± 16 %) and Actinobacteriota (Actinomycetota, 3 % ± 3 %). The other 35 phyla contributed each less than 2 % ± 1 % to the remaining 5 % ± 6 % diversity of the air microbiome (Fig. 1C, Fig. S3&4, Table S1, Dataset S2). Surface ocean communities were similarly dominated by Proteobacteria (39 %±10 %), Bacteroidota (35 %±11 %) and Cyanobacteria (15 %±11 %). The other 20 phyla contributed each less than 2 %±3 % to the remaining 11 %±8 % diversity (Fig. 1C, Fig. S3&4, Table S1, Dataset S3).

Proteobacteria and Actinobacteriota had high RSAs and high prevalences (proportion of samples in which a lineage occurs) in both biomes, yet certain lineages within these phyla were more abundant in either biome. For example, the genus *Pseudomonas* had a high prevalence (P: 67 %) and mean RSA (4 % ± 1 %) in the air, but was rare in the surface ocean (P: 17 %, RSA: 0.004 % ± 0.01 %), indicating a terrestrial source or association with the air microbiome (Fig. S5&S6). *Pseudomonas* has previously been shown to be associated with air microbiomes of the marine boundary layer (10), as well as urban, rural, mountainous, coastal (36), and forest environments (37), to name a few. Several *Pseudomonas* species are capable to initiate droplet freezing at relatively high temperatures (>−7 °C), either by acting as ice nucleating particle or by the expression of ice-nucleating proteins (24; 38).

Firmicutes were significantly more abundant and common in air (P: 100 %, RSA: 24 % ± 31 %) compared to the surface ocean (P: 47 %, RSA: 0.005 % ± 0.01 %, Fig. 1C, paired Wilcoxon signed rank test: *p*<0.0001, Dataset S4). Of the Firmicutes found in the air, 69 % ± 27 % affiliated with endospore-forming Bacilli (Dataset S6). The ability to form endospores could offer this clade an advantage in aerial dispersal, enhancing its resistance to atmospheric stressors such as UV radiation, desiccation, and limited nutrient availability (39). Bacteroidota were present in all air and ocean samples, with RSAs in the air being significantly lower (18 %±19 %) as compared to the surface ocean (35 % ± 11 %, Fig. 1C, paired Wilcoxon signed rank test: *p*<0.0001, Dataset S4).

Overall, Cyanobacteria RSA was significantly lower in the air (11 % ± 14 %) compared to the surface ocean (15 % ± 11 %, Wilcoxon signed rank test: *p*<0.01), however, when comparing pairs of air-water samples, the statistical analyses fail to find significant differences between air and ocean (paired Wilcoxon signed rank test: *p*=0.05, Dataset S4). Cyanobacteria found in air and water mainly belonged to *Prochlorococcus* and *Synechococcus*, highly abundant marine genera responsible for a substantial part of marine primary productivity (40; 41). While *Synechococcus* was similarly abundant in the air and ocean microbiomes (7 % ± 9 % and 8 % ± 14 %, respectively), *Prochlorococcus* was significantly less abundant in the air (1 % ± 2 %, Dataset S4) as compared to the surface ocean community (6 % ± 10 %, Wilcoxon signed rank test: *p*<0.001). The RSAs of *Prochlorococcus* and *Synechococcus*, the third and fourth most abundant genus level clades in the surface ocean, followed a contrasting pattern across latitudes (Fig. S6). We did not observe previously reported increasing *Prochlorococcus* abundances in the surface ocean microbiome towards the equator (42). In agreement with an expected northwards shift of increased *Prochlorococcus* abundances, due to global warming related shifts of species niches (41) and a northward expansion of the oligotrophic subtropical gyre (43), we observed a peak in *Prochlorococcus* RSA between 47° N and 35° N (Fig. S6). This is 7° further North than the suggested peak abundance of *Prochlorococcus* between 40° N and 40° S under recent (1970-2000) climate conditions. However, the observed RSAs might be impacted by seasonal patterns (44, Fig. S1).

Acidobacteriota, Gemmatimonadota and Deinococcota showed high prevalences in the air (69 %, 64 % and 50 %) with RSAs of 1.1 % ± 1.7 %, 0.4 % ± 0.7 % and 0.4 % ± 1.1 % but were absent in the surface ocean. These clades are often described for soil environments, again, suggesting terrestrial sources for the air microbiome (45; 46). Deinococcota are known for lineages with exceptional resistance towards desiccation, UV radiation and other environmental stressors (47). This resilience could increase their likelihood to survive exposure to the air environment.

Marinimicrobia (SAR406) are abundant in mesopelagic (48) and in mesotrophic oceanic regions, in particular in oxygen minimum zones (49). 39 % of air samples contained Marinimicrobia with generally low RSAs of 0.1 % ± 0.3 % (Fig. S3&S4, Dataset S2&S3). We found Marinimicrobia in all water samples, increasing in RSA towards the equator (Fig. S4).

SAR11 (Pelagibacterales) was the second most abundant genus level clade of the surface ocean, however, SAR11 RSAs were overall low compared to previously described relative cell abundances of 30 to 40 % in the Longhurst provinces WTRA, NATR, and NADR (42, Fig. S6, primer coverage 92 %). We observed a peak SAR11 RSA of 37 % at 26° N, and an average RSA of 12.8 % ± 8.6 %. The mismatch between SAR11 RSA and cell abundances might be due to the clades low 16S rRNA gene copy numbers (50).

Throughout the transect, archaeal prevalence and RSA were significantly lower in the air (P: 39 %, RSA: 0.07 ± % 0.15 %) compared to the surface ocean (P: 100 %, RSA: 1.3 % ± 1.5 %, Wilcoxon signed rank test: *p*<0.0001, Dataset S4). Archaeal RSAs and cell abundances in the air have been reported to be 1 to 6 orders of magnitude lower than bacterial RSAs (51; 52). Also, the low archaeal RSAs in the surface ocean align with previous findings, which report archaeal abundances in surface ocean waters ranging from 1 % to 10 %, with increasing relative abundances with depth (53).

### Microbial diversity of the atmosphere and surface ocean in the North East Atlantic Ocean increase towards the equator

Latitudinal diversity gradients are a widely recognized ecological pattern (54). For surface ocean microbial communities, an increase in diversity from the poles toward the equator, and a peak in diversity at intermediate latitudes have been observed (55; 56; 57), with temperature likely being a major driver (58). In general, microbial latitudinal gradients are considered to be weaker than those of larger organisms, potentially due to high abundances, high diversification rates, low extinction rates and effective long-range dispersal of microorganisms (55; 58).

Surface ocean microbial communities showed a strong negative correlation of richness and Shannon entropy with latitude (Fig. 2C, Fig. S11, Dataset S15), i.e., diversity increased towards the equator (Spearman rank correlation coefficient *ρ* = − 0.79 and *ρ* = − 0.75, respectively, *p* < 0.0001. Dataset S4). Inverse Simpson diversity showed a similar, but weaker correlation (*ρ* = − 0.64, *p* < 0.0001, Dataset S4&S15, Fig. S11). Airborne communities showed a similar though less pronounced latitudinal diversity gradient (Dataset S4&S15, Fig. S7&S11). Thus, diversity in both biomes, air and water, increased towards the equator (Fig. 2A-E) supporting studies that find peak microbial diversity in equatorial waters (56; 58), and suggesting similar trends for the air above.

**Figure 2.**
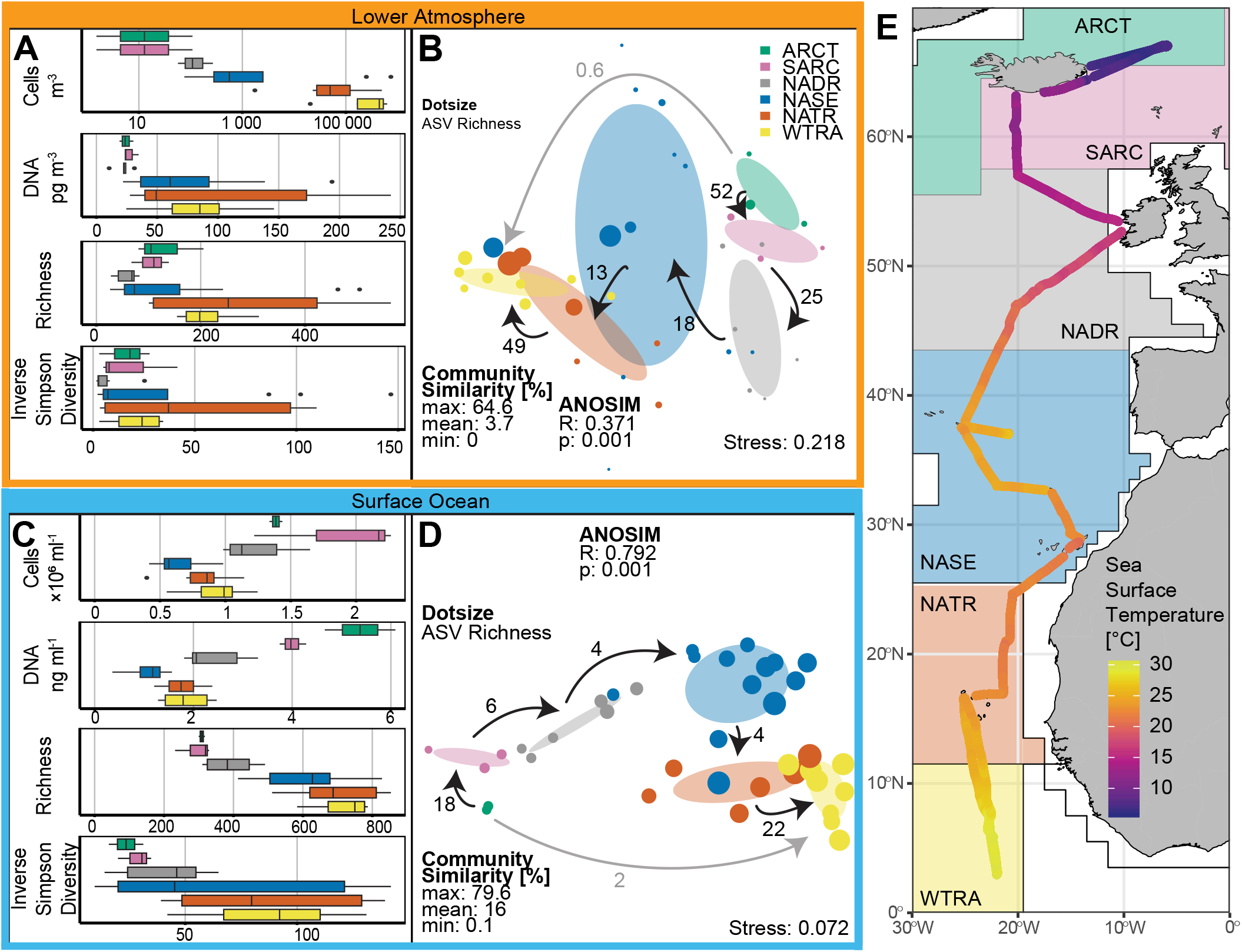
Changes in microbial alpha and beta diversity in atmospheric and oceanic communities along the latitudinal transect, grouped by oceanic provinces (33). A: Cell abundance and DNA concentration per m^3^ air (∼3.5 m above the ocean surface), community richness and inverse Simpson diversity in the sampled provinces. Green: Atlantic Arctic (ARCT), purple: Atlantic sub-Arctic (SARC), grey: North Atlantic Drift (NADR), blue: North Atlantic subtropical gyre (NASE), orange: North Atlantic Tropical Gyral (NATR), yellow: Western Tropical Atlantic (WTRA). B&D: NMDS of air (B) and surface ocean (D) microbial communities, colored by oceanic provinces. Black arrows indicate North to South gradient. Numbers at arrows show the percentage of shared ASVs between each province (corrected to group size). Grey arrow shows change from northernmost (ARCT) to southernmost (WTRA) province and the percentage of shared ASVs between the two provinces. C: Cell abundance and DNA concentration per mL surface ocean water (∼3 m depth) and the associated community richness and inverse Simpson diversity. DNA: 1 outlier in WTRA not shown (16.7 ng mL^−1^ DNA, sample ES21C08_08w). E: Sea surface temperature along the transect with oceanic provinces.

Surface ocean microbial richness correlated with sea surface temperature and dissolved oxygen concentration (Fig. 2E, Dataset S4&S15 for *ρ*, and *p*-values). We observed a weak correlation of richness with chlorophyll-*a* fluorescence, cell abundance, and salinity (Dataset S4). Similar to the water communities, air community richness showed a correlation with sea surface temperature and ambient air temperature as well as surface ocean dissolved oxygen concentration. Surface ocean chlorophyll-*a* and salinity did not correlate with the air microbial community richness (Dataset S4), suggesting that additional environmental factors impact the latitudinal diversity of the air community.

Cell abundances in the air above the North East Atlantic Ocean increased towards the equator, together with increasing concentrations of extractable DNA (Fig. 2A, Fig. S11). Given the Coriolis µ has a D_50_ < 0.5 µm (diameter at which at least 50 % of particles are collected), cell numbers are likely underestimated, however, the observed trends are expected to remain consistent. Cell abundances ranged from zero (not detectable) to 0.74 × 10^6^ cells m^−3^ (0.12 × 10^6^ ± 0.21 × 10^6^ cells m^−3^, median: 500 cells m^−3^, Table S7). Cell abundances observed in the northern North Atlantic (ARCT, SARC, NADR, 55 ± 79 cells m^−3^) are in accordance with previous reports (5). However, cell abundances greatly increased towards the South, spanning five orders of magnitude (Fig. 2A, Fig. S8&S11). The high cell abundances of 0.3 × 10^6^ ± 0.2 × 10^6^ cells m^−3^ close to the equator (NATR, WTRA) coincided with land-derived air masses and increased particle number concentrations of 502 ± 174 particles cm^−3^ in the air (Fig. 3A,B, Fig. S7). Similar cell abundances were previously described for airborne prokaryotes over the Red Sea (7). However, high particle concentrations at northern latitudes did not coincide with increased cell abundances, possibly due to differing aerosol compositions. The increase in alpha diversity and cell abundance in the tropics might be caused by air masses originating from Africa (Fig. 3A, Dataset S14), adding microbial lineages and cells. It has to be noted here that the sampling height of 3.5 m was comparatively close to the ocean surface. Nevertheless, long-range transported aerosols typically also reach the lower sampling heights in the convectively mixed marine boundary layer. In addition, increased nutrient availability (e.g., phosphorous, iron) in the air masses due to long-range transport (26), together with increased temperatures and humidity may render the air itself more habitable. Estimates of bacterial cell abundances over the open oceans currently used to model atmospheric processes (1 × 10^4^ cells m^−3^, range: 1 × 10^1^ – 8 × 10^4^ cells m^−3^, 3) are up to one order of magnitude lower than our findings. This discrepancy may be due to an underestimation of long-range transport of land-derived organisms and should be considered in future studies and models.

**Figure 3.**
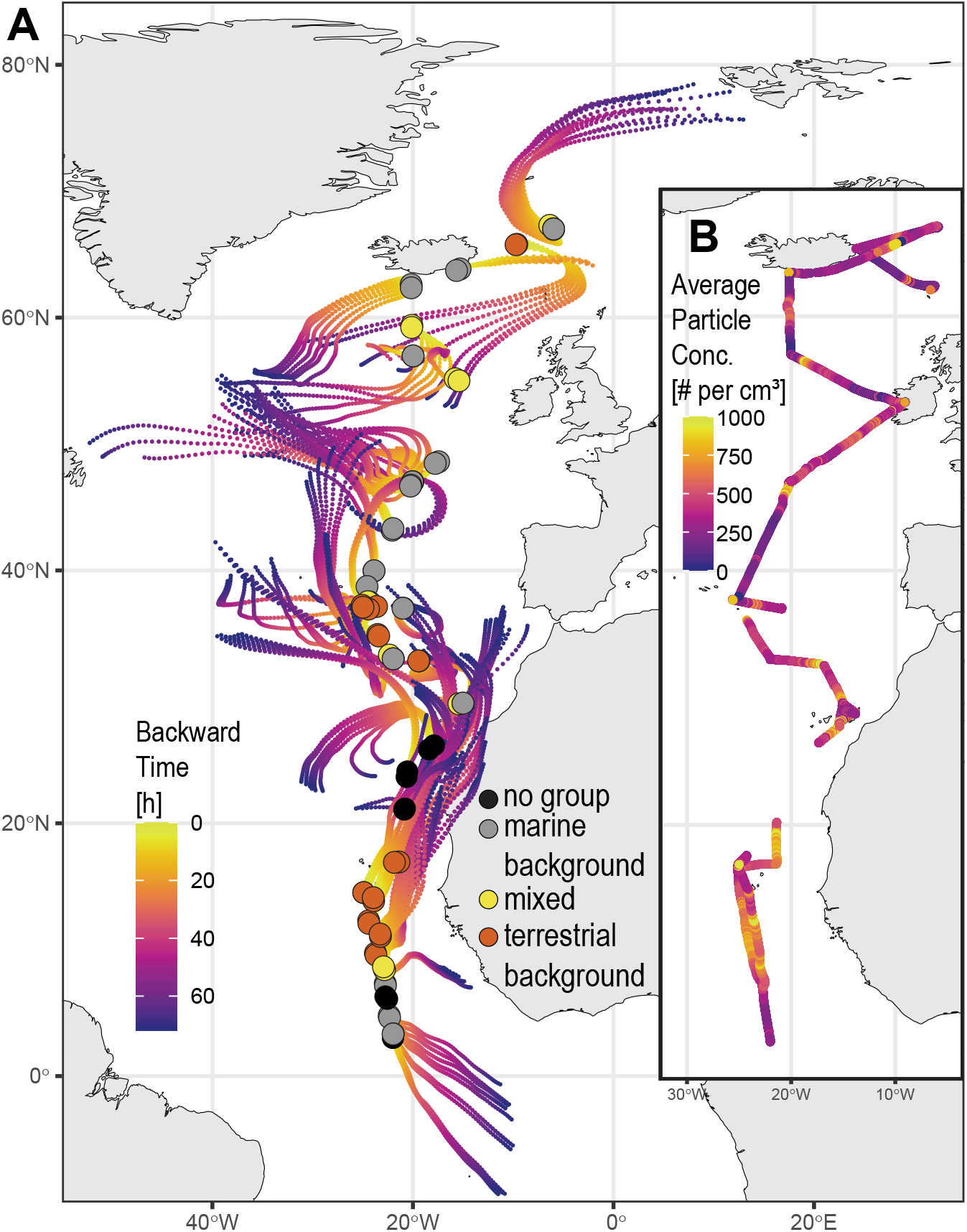
Sampled air masses. A: HYSPLIT three-day backward trajectories during sampling times. The color gradient shows the time until air masses reached the sampling location (up to 72 h). Colored circles represent different air mass origins (see Methods). Samples not assigned to a group due to unavailable data are shown as black circles. B: Total aerosol particle number concentration per cm^3^ along the cruise track.

Surface ocean cell abundances showed a bimodal distribution with maximum cell abundances at subpolar (1.49 × 10^6^ ± 0.5 × 10^6^ cells mL^−1^; SARC, ARCT, 60° N – 67° N) and equatorial latitudes (0.87 × 10^6^ ± 0.3 × 10^6^ cells mL^−1^; WTRA, 10° N – 3° N) and minimal cell abundances in the subtropical gyre at around 30° N (NASE, 0.42 × 10^6^ ± 0.14 × 10^6^ cells mL^−1^, 35 °N – 25° N, Fig. 2C, Fig. S8, Fig. S11). We also observed a clear geographic clustering by the sampled Longhurst provinces (33), in both air and ocean communities, with very little diversity overlap between the provinces in both biomes (Fig. 2B&D). Similar spatial separation was observed along the same transect in the surface ocean’s physical and photophysiological characteristics of the autotrophic community (59). This suggests that despite the strong differences between the air and ocean biomes, strong local processes and environmental pressures must occur and shape the communities of both biomes alike. To our knowledge, this is the first time a latitudinal microbial diversity and cell abundance gradient is observed in the atmosphere, highlighting the possible influence of terrestrial and marine sources on the air community structure.

### Air mass origin and aerosol concentration impact surface ocean and air microbial communities

To further understand the influence of the air mass origin on the microbial communities present at our sites, we studied whether microbial community structure correlated with i) the distance of the sampling site to the nearest land mass (distance to shore), ii) the length of the air masses travelled within 3 days prior to their arrival at the sampling site (trajectory length), and iii) the potential influence of land-derived air masses (land influence). For the latter we used the fraction of backward trajectories (BTs) crossing land masses within the past 3 days divided by the total number of calculated BTs (usually six) for each individual sample (Fig. S7B&F, see Methods section), together with locally observed aerosol number concentrations, as well as the observed particle number size distributions (Fig. S9E).

The microbial community richness in water and air did not correlate with distance to shore (i, Dataset S4). However, in the air, some lineages typically associated with soil or freshwater environments, e.g., Acidobacteriota and Gemmatimonadota (45; 46), showed decreasing RSAs with increasing distance from shore. This suggests a clade-specific decrease of RSA with increasing distance to land, while the overall community composition stays relatively unaltered. Both, ocean and air microbial richness increased with trajectory length, likely due to the air masses increased potential exposure to different terrestrial and marine ecosystems (ii, Fig. S7A&E).

Using land influence (Fig. 3A, Fig. S7B&F, Dataset S14) and particle concentrations (iii, Fig. 3B, Fig. S7C&G), we grouped the sampling sites into three categories. Marine background sites (n=16) are sites were BTs did not traverse terrestrial areas and the median particle concentration was lower than 400 particles per cm^3^. At terrestrial background sites (n=11), at least one BT crossed terrestrial areas and the median particle concentration exceeded 400 particles per cm^3^. The remaining sites (n=6) are considered mixed background (marine or terrestrial origin, Fig. 4, Fig. S9, see Methods section for details).

**Figure 4.**
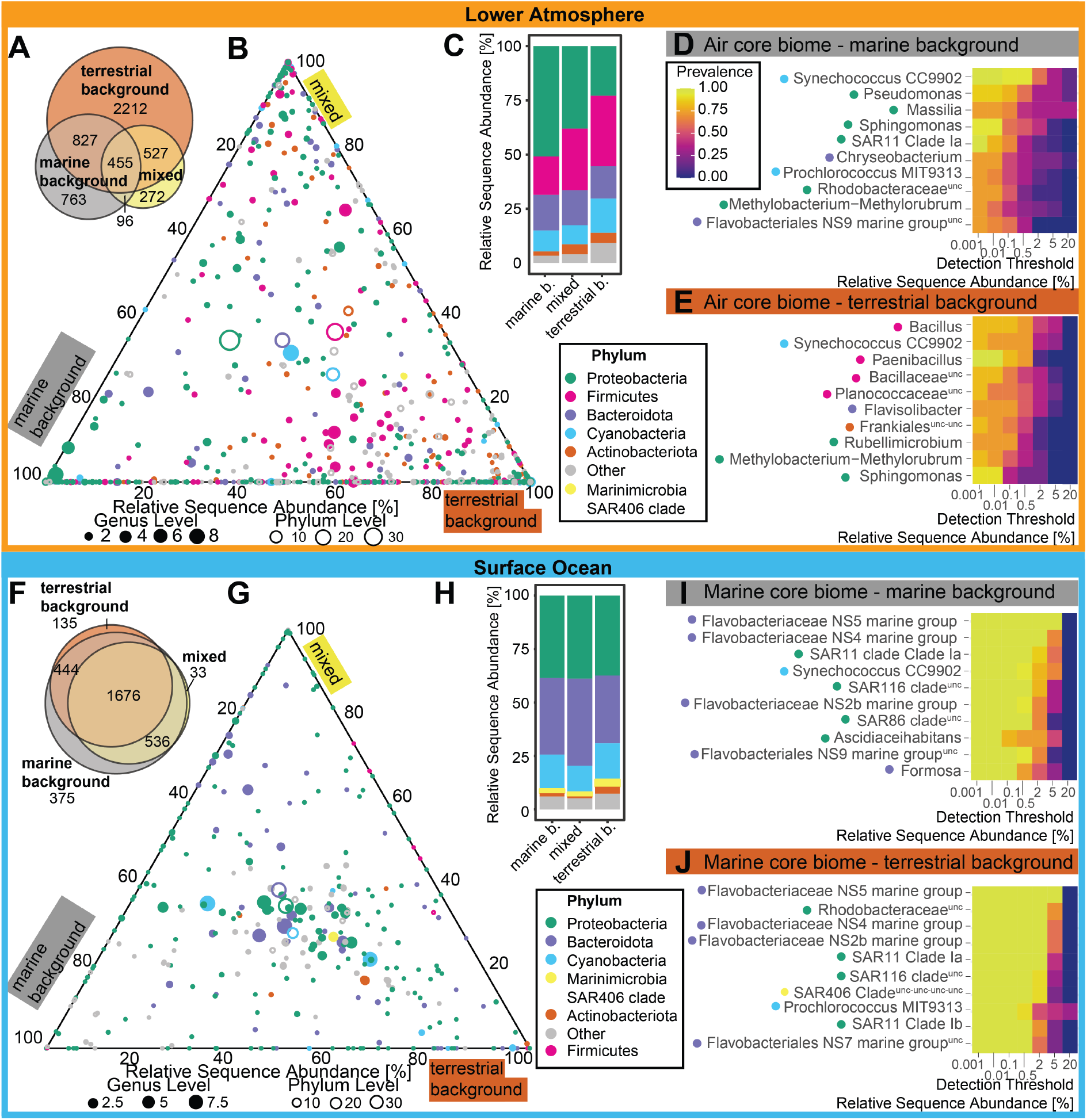
Air and surface ocean community composition under terrestrial, marine and mixed conditions. A&F: Scaled Venn diagrams (corrected for group sizes) showing number of observed 16S rRNA gene sequence amplicon variants (ASVs). B&G: Ternary plots. Closed circles represent taxonomy at genus level, open circles represent taxonomy at phylum level, circle size represents relative sequence abundance of the respective taxon averaged across all samples of the respective biome. Dot position indicates the distribution across the conditions. Top 5 abundant phyla of either biome are distinct colors, while the remaining phyla (“Other”) are grey. C&H: Relative phylum level sequence abundance per group. D, E, I & J: Genus level core microbiomes of air and ocean communities considering different air mass histories. Top 10 genus level clades are shown and their prevalence (i.e., proportion of samples the genus occurs in) is color-coded (legend in plot D) for each relative sequence abundance threshold.

Air and surface ocean microbial alpha diversity were elevated under terrestrial background conditions, independent of latitude (Fig. 3, Fig. S9&S7). Terrestrial-derived microbes transported within these air masses likely increase the local diversity (8; 60; 61). The associated change in the availability of trace elements, such as soluble iron and silicon through dust input to the surface ocean, might cause or enhance the observed community shifts (62).

#### Air microbiome

Airborne microbial communities showed distinct community structures depending on the air mass background (Fig. 4A-E, Fig. S9B&S14), characterized by different core microbiomes. We define the core microbiome as the most relevant microbial lineages, determined by their prevalence and relative sequence abundance (see methods section). Air microbial communities with a terrestrial background harbored almost twice as many ASVs as air communities with a marine background (Fig. 4A). While cyanobacterial RSA was similar at stations of all three categories (Fig. 4B,C, Fig. S14), the RSA of Proteobacteria in the air was significantly higher at sites with a marine background compared to those with a terrestrial background (Wilcoxon rank sum test: *p*<0.05). A terrestrial background led to an air core microbiome dominated by Firmicutes lineages (Fig. 4E), a clade known to be highly abundant during dust influence (63). Also, the order Frankiales (Actinobacteriota), known to be associated with North African dust (64) was found in the air core microbiome during terrestrial background conditions (Fig. 4D&E). A marine background led to an air core microbiome dominated by marine lineages such as *Synechococcus, Prochlorococcus* (41) and SAR11 (42), highlighting the importance of surface ocean sources for the air microbiome during marine background conditions (Fig. 4D). Interestingly, we also find *Synechococcus* at high RSAs in the air microbiome of apparent terrestrial origin, highlighting the clade’s importance in the marine boundary layer, and indicating a high aerosolization potential. Organisms affiliating with *Pseudomonas, Massilia*, and *Sphingomonas* were also members of the air core microbiome during marine background conditions, which may reflect the widespread distribution of these genera in diverse global ecosystems (65; 66; 67).

#### Surface ocean microbiome

The surface ocean microbial communities, as compared to the air communities, showed very different diversity patterns in response to air mass backgrounds (Fig. 4F-J). While the microbial richness per ocean sample increased under terrestrial background influence (Fig. S9C), total richness within each category stayed very similar (Fig. 4F). RSAs of certain bacterial phyla, e.g., Marinimicrobia (SAR406), Actinobacteriota and Firmicutes significantly increased (Wilcoxon rank sum test: *p*<0.05, <0.01, <0.01, respectively) during terrestrial background conditions (Fig. 4H). In contrast to the air community, genus level clades were very similarly distributed between the three conditions (Fig. 4G). The genus level core microbiomes during terrestrial and marine background conditions mainly consisted of typical marine clades such as Flavobacteriaceae groups, SAR11 and Cyanobacteriaceae genera (Fig. 4I&J, 41; 42). We propose a minor shift in surface ocean community structure due to the introduction of terrestrial-derived clades. Small changes may be overshadowed by high microbial abundances, the more homogeneous nature of the surface ocean community, and the influence of surface ocean physical and biological dynamics (59).

### Exchange of surface ocean and air communities may be lineage-selective

Several environmental processes can influence the air-sea exchange of particles and gases. Higher wind speeds lead to an increased mixing of the surface ocean (68), increased wave formation, bubble bursting and enhanced sea-spray aerosol formation (69). Hence, it is expected that higher wind speeds would be associated with higher microbial exchange rates. While we could observe a negative correlation between the Bray-Curtis dissimilarities and wind speed, i.e., communities tended to be more similar with higher windspeeds, this trend was not significant (*R*= − 0.35, *p*=0.062) and requires further investigation. Low wind speeds during sample collection (5.2 ± 2.3 m s^−1^) may have limited the exchange between surface ocean and air communities (high Bray-Curtis dissimilarities: 0.74 to 1), yet are representative for the global average wind speed of 6.6 m s^−1^ (70). It remains unclear whether changes in microbial air communities above the oceans may thus be caused by microbes introduced through long-range transport, and how air mass origin, locally aerosolized microbial organisms, and wind speed shape the air microbiome.

We investigated whether certain microbial lineages are predominantly associated with surface waters or air masses, and which lineages are potentially exchanged between the two biomes. To do this, we applied a rule-set considering prevalences, RSAs (using pairwise Wilcoxon signed rank tests) and correlations of the RSAs of each lineage between the two biomes (Spearman’s rank correlation test, two-sided). The classification of lineages is summarized in Fig. 5 and described in Table S6 and the SI Appendix Methods (Dataset S5).

**Figure 5.**
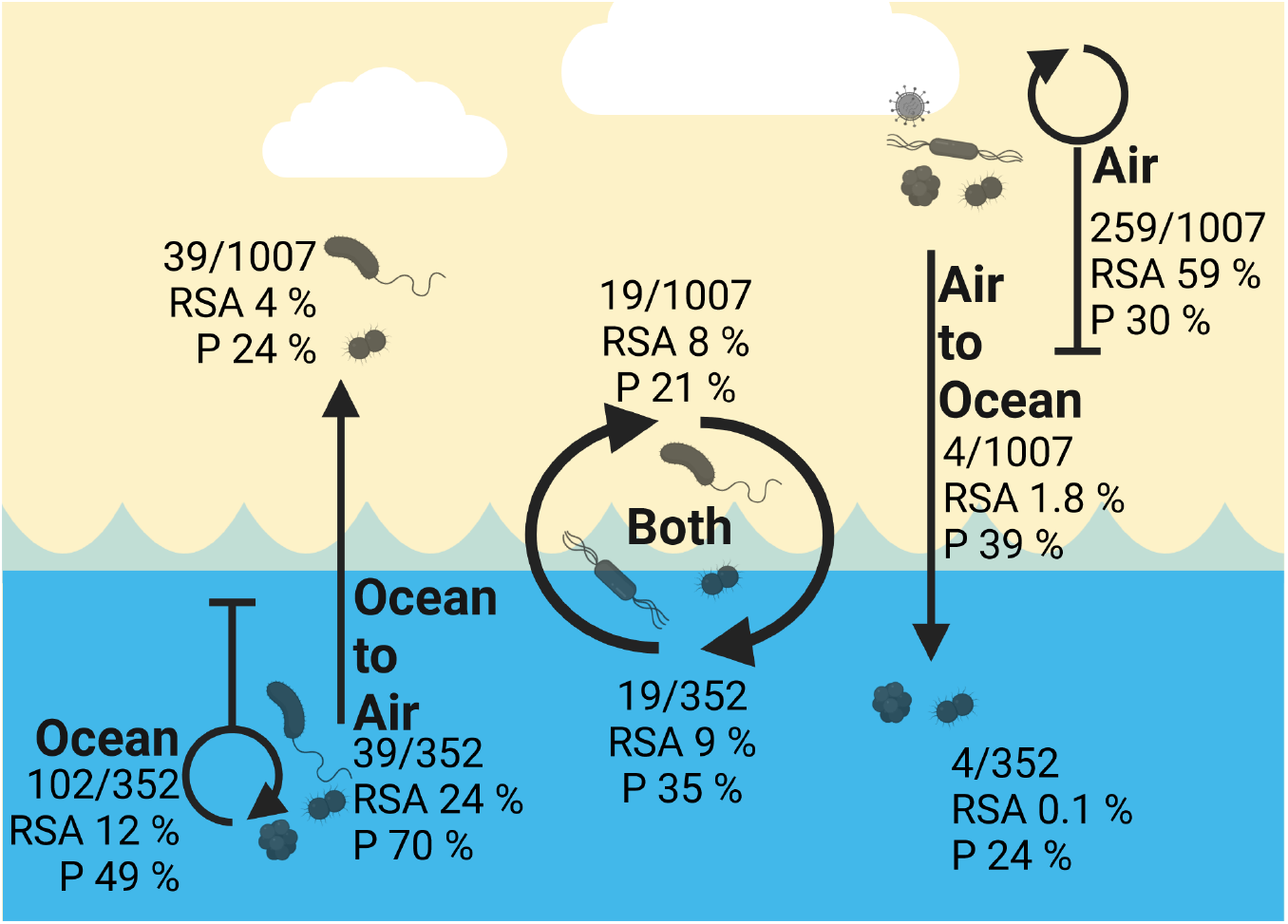
Schematic exchange of microbial lineages between the lower atmosphere and the surface ocean. Ocean to Air: Ocean associated, impacting the air microbiome, Ocean: Ocean associated, not impacting the air microbiome. Both: potentially associated with both biomes, indicating a high degree of exchange. Air: Atmosphere associated, not impacting the surface ocean microbiome. Air to Ocean: Atmosphere associated, impacting the surface ocean microbiome. Numbers show genus level lineages (or unclassified higher taxonomic levels) assigned to each group, together with the total relative sequence abundance (RSA) of those lineages and the average prevalence (P) in the respective biome. Lineages which could not be assigned to one group were excluded but are listed in Dataset S5.

#### Introduction of terrestrial-derived biodiversity to the surface ocean

Of the 40 phyla found in the lower atmosphere, only Firmicutes was found to potentially impact the surface ocean community (Group “Air to Ocean”). Despite the overall low RSAs of Firmicutes in the surface ocean, high Firmicutes RSAs in the air coincided with increased RSAs in the surface ocean (Fig. 5, Fig. S4). This suggests an atmospheric source for Firmicutes found in the surface ocean, corroborating trends seen in the North Pacific (71). Firmicutes transcriptomes were isolated from sediment samples and the water column (72; 73) suggesting a potential ecological niche within the ocean realm for this clade.

Seven phyla, including Acidobacteriota, Deinococcota and Gemmatimonadota, abundant in the air microbiome (RSA up to 7 %), did not occur in any of the ocean samples (Group “Air”, Dataset S2,3&6). Even when cell abundances in the air were high, these phyla did not impact the surface ocean community. This suggests the lack of an ecological niche in the oceans for these lineages, which are commonly found in high abundances in soil (74; 75) or air (47).

We suggest that a successful implementation of airborne organisms to the surface ocean relies on high cell abundances together with high RSAs in the air. However, additional factors, e.g., osmotic stress tolerance, cell surface composition, nutrient, carbon and energy availability, and the length of exposure, likely influence the inmixing and survival of air-derived microbial cells in the surface ocean (76).

#### Introduction of ocean-derived biodiversity to the atmosphere

Three of the 23 phyla found in the surface ocean showed an impact on the air community; Thermoplasmatota, Chloroflexi (Chloroflexota) and Desulfobacterota (Fig. S4). At genus level, 39 of the 352 genera found in the surface ocean (e.g., *Prochlorococcus*) were abundant in the air, indicating they are easily aerosolized (Group “Ocean to Air”, Fig. 5), supporting previous findings from sites in the North Pacific (71). In fact, *Synechococcus* RSAs were almost identical in both biomes (Group “Both”), suggesting efficient aerosolization of this typical marine clade (Fig. S6, Dataset S5).

Four phyla were found predominantly in the ocean, Dadabacteria (Dadabacteriota), Margulisbacteria (Margulisiibacteriota), SAR324 marine group B and PAUC34f (Fig. S4), and almost a third of the genera found in the surface ocean were not found in the air, suggesting a non-random aerosolization/non-aerosolization or a lack of the ability to stay airborne or survive in the air.

Actinobacteriota were present in all air and surface ocean samples and showed similar RSAs in both biomes (Fig. 1C). Actinobacteriota RSAs increased during terrestrial background conditions (Fig. 4, Fig. S14) in air and water. However, while the surface ocean was dominated by typical aquatic lineages belonging to the class of Acidimicrobiia (77), the lower atmosphere was dominated by lineages belonging to the class of Actinobacteria (Actionmycetes, Fig. S6, Dataset S5).

RSAs of SAR11 genera did not correlate between ocean and air communities while showing high abundances in both environments (Fig. S6, Dataset S5). Locally aerosolized SAR11 likely accounted for the majority of airborne SAR11 due to the sampler’s proximity to the sea surface (3.5 m), yet their small cell size and surface properties (78) might also facilitate long-range transport. The ability to remain airborne over extended periods could contribute to the global distribution and abundance of SAR11 (79). It is important to note that the used air sampler is limited in capturing cells smaller than 0.5 µm, which may lead to an underestimation of SAR11 cells (80; 81) in air samples. Therefore, comparisons between air and ocean microbiomes should be approached with caution regarding lineages characterized by small cell sizes.

The observed patterns in microbial community structure in air and ocean microbiomes suggest lineage-selective aerosolization of microorganisms across the ocean-air-interface. Although further direct evidence via tracer studies and temporal tracking is needed, the observed exchange of microbes indicates highly specific biochemical or biophysical processes at the sea-air interface, with cell size, surface structure (76; 78; 82) and surface microlayer enrichment (83) being potential influencing factors.

## Conclusion

Our study provides robust community analyses across biomes and latitudes suggesting that microbial cells are aerosolized and deposited non-randomly, and that certain lineages, e.g., *Synechococcus*, are exchanged across the sea-air interface more efficiently and frequently than others. *Synechococcus* blooms may thus have a more pronounced impact on sea spray aerosol compositions and lower atmosphere carbon fluxes than other lineages, e.g., Marinimicrobia. The high relative sequence abundance of SAR11 in the atmosphere is intriguing and further work is needed to determine whether this clade, and others, are viable and metabolically active in aerial environments above the ocean. We demonstrate that airborne microbes show a latitudinal diversity gradient similar to plants and animals, indicating that atmospheric microbial communities follow ecological rules similar to those of the terrestrial and marine biosphere. Furthermore, we show that air masses with high particle number concentrations, potentially influenced by continental ecosystems, carry a higher microbial diversity. Although apparently not affecting the marine community at large, the introduced microbes and terrestrial-derived nutrients harbor the potential to alter surface ocean microbial diversity by impacting individual microbial populations. For instance, we observed substantial differences in the relative sequence abundance of Marinimicrobia, Actinobacteriota, and Firmicutes in response to changes in air mass origin. To understand the interplay of global biogeochemical cycles and climate, it is critical to further explore and directly quantify the microbial contributions to sea spray aerosol formation, marine aerosol-cloud interactions, and the hydrological cycle.

## Methods

### Sampling area and platform

Sampling and continuous measurements of the North East Atlantic Ocean, from 67° N to 3° N, roughly following the 20° W meridian, were performed on the 72-foot research vessel *S/Y Eugen Seibold* between June 2020 and May 2021, probing six oceanic provinces (Fig. S1, Table S2). A cruise track is publicly available at PANGAEA. A list of the collected samples, their locations, sample volumes, DNA concentrations, sequencing results, and blanks samples, is provided in Dataset S1 (sample list). Information on additional sampled and measured parameters can be found elsewhere (32).

### Air sampling

Air samples for microbial analyses were taken using a liquid-based cyclonic air collector (Coriolis µ, Bertin Technologies, D_50_ < 0.5 µm) located on top of the wheelhouse of the research vessel at ∼3.5 m above sea level. Samples were taken with an air-flow of 300 L min^-1^ over a time of 360 min, with a few exceptions (Dataset S1), in 15 mL collection liquid (1x phosphate-buffered saline). Liquid level was kept constant at 15 mL by adding ultrapure water (MQ; resistivity 18.2 MΩ·cm). If necessary, the final volume was adjusted to 15 mL with MQ and the collection cone vortexed shortly to resuspend particles. Strict hygiene protocols were followed and samples processed immediately after collection (for detailed protocols see SI Methods). For DNA analyses, an aliquot of 12 mL was filtered on 0.2 µm pore size filters (Isopore PC membrane filters, GTTP02500), filters were stored in 2 mL centrifuge tubes and kept at -80 °C. For microbial cell enumeration, an aliquot of 1.5 mL or 2 mL of sample was fixed with formaldehyde (f.c. 1 %) for 1 h at room temperature or at 4 °C for approximately 18 h (Dataset S1). Fixed samples were filtered on 0.2 µm, 25 mm polycarbonate membranes (Isopore™ GTTP02500, with 0.45 µm pore size cellulose acetate filters as support). Filters were stored at -20 °C.

### Surface ocean water sampling

Surface water samples were collected directly from the keel inlet using the OceanPack Ferrybox (3.2 m water depth, details below) or from Niskin bottles. When reused or not delivered sterile, sampling equipment was cleaned with 70 % ethanol and MQ. For surface water DNA samples, bottles were rinsed three times with 50 mL freshly collected sample water before the bottles were filled to 1 L. When several samples (i.e., from different depths) were taken simultaneously, samples were stored at 4 °C until filtration. Samples were filtered sequentially through 10, 3 and 0.2 µm pore size filters (Dataset S1) and stored at -80 °C. Samples for microbial cell enumeration (19 mL sample volume) were fixed similar to air samples and filtered on 0.2 µm pore size, 47 mm poly-carbonate membranes (Isopore™ GTTP04700, with 0.45 µm pore size cellulose acetate filters as support). Filters were stored as flat as possible in Analyslides (Pall, PI-BE-7231) at -20 °C. For blank samples, 100 mL of MQ was filtered for DNA, and similar volumes as sample volumes for the other sample types.

### Atmospheric and meteorological measurements and analyses

Air for continuous measurements of particle concentrations is sampled through an electrically conductive (carbon-primed) polyurethane tube with an inner diameter of 3.9 mm through a 10 m long inlet-line, located behind the main sail, resulting in a sampling height of ∼13 m above sea level. Flow rates have been optimized for low residence time while ensuring low particle losses. Total particle concentrations were measured with a condensation particle counter (WCPC, TSI Inc., Shoreview, USA, Model 3787). The median particle concentration for each sample was calculated and used to group samples based on potential air mass origins (Dataset S1). Particle number size distributions were recorded using a scanning mobility particle sizer (SMPS, TSI Inc., with electrostatic classifier model 3938 and CPC model 3787). Particle transmission efficiencies were calculated (84). For our setup and flows, the lower D_50, CPC_ was 8 nm, and the upper D_50, CPC_ was 3.6 µm (Fig. S13A, SI Dataset S16). The lower D_50, SMPS_ was 7.4 nm, and the upper D_50, SMPS_ was 3.6 µm (Fig. S13B, SI Dataset S16, smallest diameter recorded: 9.32 nm, largest: 429.4 nm). Further details on the inlet and aerosol setup can be found in Schiebel et al., 2024 (32).

### Surface ocean chlorophyll-*a*, SST, salinity, dissolved oxygen

Surface ocean chlorophyll-*a* fluorescence, SST, Salinity and dissolved oxygen were measured with a *FerryBox* (*OceanPack*, SubCtech GmbH) system, continuously sampling water from ∼3 m water depth. A detailed description and data can be found in (32; 59; 85). Values during the sampling times were averaged and are summarized in Dataset S1.

### Backward trajectory calculation

1. day backward trajectories were modelled using HYSPLIT (Hybrid Single-Particle Lagrangian Integrated Trajectory) for every hour starting at the current location of the boat along the cruise track with the starting altitude fixed at 200 m above sea level (86; 87) with the meteorological dataset GDAS1 (1° by 1° resolution). The respective backward trajectories for each air sample were extracted to calculate the percentage of trajectories traversing land masses within the past 3 days after cutting them in case moderate rain occurred (more than 0.5 mm h^−1^) to account for particle loss due to wet deposition. Averaged values of data products (relative humidity, height above ground, length, pressure, potential temperature, rainfall, air temperature, mixing depth) are included in Dataset S1.

### DNA extraction

Due to low biomass in air samples, different DNA extraction protocols were tested and modified to maximize DNA yield and integrity. Different extraction protocols and commercially available DNA extraction kits were tested (Dataset S7). Samples were finally extracted using the ChargeSwitch™ Forensic DNA Purification Kit (ThermoFisher Scientific), which proved suitable for air and water samples, with slight modifications (freeze-thaw cycles, additional lysozyme digest, for details see SI Methods). DNA eluates were stored at -70 °C until sequencing.

### Library preparation and sequencing

Negative and positive controls were sequenced together with the samples. Library preparation and sequencing was conducted by StarSEQ (StarSEQ GmbH, Mainz, DE). The v4-v5 region of the 16S rRNA gene was sequenced using a dual-index strategy based on the protocol with minor modifications (88). Amplicons were generated in a single step of 34 cycles using the primer combination 515F (5-GTGNCAGCMGCCGCGGTA-3, 89) and 924R (5-CCCCGYCAATTCMTTTRAGT-3, 90). The final library was sequenced on the Illumina MiSeq platform (MiSeq™, Illumina, San Diego, CA, USA) in paired-end mode at a respective length of 300 nucleotides. Raw reads were de-multiplexed and adapters were trimmed. Primers were tested on their coverage using TestPrime 1.0 (Dataset S8, 91). Sequencing data is publicly available through PRJNA1148129 (data is uploaded and will be released upon publication at NCBI).

### Staining of microbial cells

A circular section of filter samples was cut out with a sterile scalpel and placed with the sampled side in a 10 µL DAPI (4’6-diamidino-2-phenylindole, 1 µg mL^−1^) for 10 min at room temperature in darkness, followed by short washes in ultra pure water and >70 % ethanol. Filter pieces were dried and mounted in Citifluor:Vectashield (4:1, v/v; Citifluor, London, UK; Vector Laboratories, Burlingame, CA, USA).

### Automated image recording and processing for cell enumeration

Images were visualized using a Zeiss AxioImager.Z2m microscope, equipped with a Zeiss Colibri 7 LED and a Multi Zeiss 62 HE filter cube. Images were recorded using a charged-coupled device (CCD) camera, the Zeiss AxioVision software and a custom-build macro (92). Images were analyzed with the Automated Cell Measuring and Enumeration tool (ACME). For details refer to SI Appendix, Methods. Representative cell pictures are compiled in Dataset S12.

#### Calculation of cell abundances in air and water

The derived numbers of cells per field of view were used to calculate average cell abundances in the air (cells m^−3^) and in the water (cells mL^−1^), for details refer to SI Appendix, Methods. Low-biomass (air samples and blanks) raw counts per collected volume were corrected before calculating final cell abundances (SI Appendix, Methods Cell enumeration).

### Bioinformatic analysis and visualization

Analyses and visualization were performed in the software environment R (93). For details refer to the publicly available workflows and the SI Methods section.

#### Illumina read processing and taxonomy assignment

The “Diverse Amplicon Denoising Algorithm” (DADA2, 94) was used to investigate read quality, to filter and trim and merge reads, calculate sample inference, remove chimeric sequences and taxonomy assignment with the SILVA 138.1 database. For details on sequence processing, including the effects of different filtering and trimming parameters refer to SI Methods section “Illumina read processing and taxonomy assignment” & Dataset S9.

#### Removal of potential contaminant sequences

The effect of several bioinformatic strategies to remove potentially contaminating sequences was investigated. DNA concentrations above the detection limit were observed for some of the field blanks. All field blanks were extracted and sequenced together with the samples. A summary and evaluation of the effects of the 47 tested bioninformatic decontamination strategies on the community structure is summarized in Dataset S10 and explained in more detail in the SI Appendix, Methods section. Combination 4 (Table S3) was considered to be the most appropriate procedure, efficiently removing contaminant sequences from samples (and blanks) while minimizing the risk of introducing unjust trends into the sample set. Here, we first used the decontam package (95) for low biomass samples and then removed the maximum occurring read count in any of the blanks from each sample, followed by the removal of potential human and reagent contaminants as listed in Dataset S11. A detailed comparison of the community structures and shifts, before and after applying different decontamination approaches, is shown in Fig. S10 and summarized in Tables S3-5.

#### Visualization of community data

Community composition analysis and visualization was performed using a custom R workflow VisuaR and accessory custom R scripts. Water and air samples were grouped based on different environmental parameters, e.g., water vs air. Samples were also assigned to different biogeographical provinces (33) using a latitudinal rule-set as defined in (59) and summarized in Table S2. Data was analyzed with and without contaminant removal (Fig. S10), and with and without transformation (Fig. S12) to ensure high-quality, reproducible results.

#### Grouping of samples based on air mass origin and particle concentration

To classify samples based on air-mass origin (marine, terrestrial, mixed/unclear), the median particle concentration during sampling together with information on 3-day backward trajectories crossing land masses before arriving at the sampling location was used. For water samples, the grouping of the corresponding air samples was used. An upper limit of 400 particles cm^−3^ for marine background conditions was set based on (96) which reported average marine particle number concentrations ∼435 particles cm^−3^ between 75° N and 0° N and well within the upper threshold of 600 particles cm^−3^ defined as clean marine conditions by (97). *Marine background:* Median particle concentration during sampling <400 particles cm^−3^ and no 3-day backward trajectory traversed land. *Terrestrial background:* Median particle concentration during sampling >400 particles cm^−3^ and at least one 3-day backward trajectory traversed land. *Mixed:* The remaining samples. To evaluate the quality of groupings of air mass origin based on particle concentration and air mass backward trajectories, particle number size distributions of the different categories were investigated (Fig. S9E, Dataset S17), showing elevated particle concentrations, in particular for smaller particles between 10 nm and 30 nm during the assumed influence of terrestrial air masses.

## Acknowledgements

Our appreciation goes to our skilled captains Kaarel Kruusmägi and Karl Vahtra for ensuring safe and productive journeys, and to the rest of the crew of the *S/Y Eugen Seibold*, to Argo Kruusmägi, Margus Zaharov and Karl Trink for their endless support, and to Janette Possul, Adeele Kuslap, Marharyta Kalashnikova and Marge Piirments. We acknowledge the work of Sonja Endres, Florian and Jeannine Ditas in the initial setup of the on-board laboratories. We thank the mechanical and electrical workshops of the MPIC for their valuable support. We acknowledge the support of the S/Y *Eugen Seibold* project by the Werner Siemens foundation and the Max Planck Society, the Max Planck Institute for Chemistry and the Max Planck Graduate Center with the Johannes Gutenberg University of Mainz (MPGC).

## Author contributions statement

According to Contributor Roles Taxonomy (CRediT, https://casrai.org/credit/):

Conceptualization: IHdA, CP, SER

Data curation: IHdA, HMA, HAS, DW

Formal analysis: IHdA

Investigation: IHdA, HMA, MLC, AD, HAS, SB, RS, AL

Software: IHdA

Supervision: CP, SER

Validation: IHdA, HMA, HAS, SB

Visualization: IHdA

Writing – original draft: IHdA, SER Writing – review & editing: All authors

## References

1. Womack, A. M., Bohannan, B. J. M. & Green, J. L. Biodiversity and biogeography of the atmosphere. Philos. Transactions Royal Soc. B: Biol. Sci. 365, 3645–3653, DOI: 10.1098/rstb.2010.0283 (2010).

2. Després, V. R. et al. Primary biological aerosol particles in the atmosphere: a review. Tellus B: Chem. Phys. Meteorol. 64, 15598, DOI: 10.3402/tellusb.v64i0.15598 (2012).

3. Burrows, S. M., Elbert, W., Lawrence, M. G. & Pöschl, U. Bacteria in the global atmosphere – Part 1: Review and synthesis of literature data for different ecosystems. Atmospheric Chem. Phys. 9, 9263–9280, DOI: 10.5194/acp-9-9263-2009 (2009).

4. Lewis, E. R. & Schwartz, S. E. Sea salt aerosol production: Mechanisms, methods, measurements and models—A critical review. In Geophysical Monograph Series, vol. 152, DOI: 10.1029/152GM01 (2004).

5. Mayol, E., Jiménez, M. A., Herndl, G. J., Duarte, C. M. & Arrieta, J. M. Resolving the abundance and air-sea fluxes of airborne microorganisms in the North Atlantic Ocean. Front. Microbiol. 5, 557, DOI: 10.3389/fmicb.2014.00557 (2014).

6. Mayol, E. et al. Long-range transport of airborne microbes over the global tropical and subtropical ocean. Nat. Commun. 8, 201, DOI: 10.1038/s41467-017-00110-9 (2017).

7. Yahya, R. Z., Arrieta, J. M., Cusack, M. & Duarte, C. M. Airborne Prokaryote and Virus Abundance Over the Red Sea. Front. Microbiol. 10, 1112, DOI: 10.3389/fmicb.2019.01112 (2019).

8. Mescioglu, E. et al. Aerosol Microbiome over the Mediterranean Sea Diversity and Abundance. Atmosphere 10, 440, DOI: 10.3390/atmos10080440 (2019).

9. Cho, B. C. & Hwang, C. Y. Prokaryotic abundance and 16S rRNA gene sequences detected in marine aerosols on the East Sea (Korea). FEMS microbiology ecology 76, 327–341, DOI: 10.1111/j.1574-6941.2011.01053.x (2011).

10. Lang-Yona, N. et al. Terrestrial and marine influence on atmospheric bacterial diversity over the north Atlantic and Pacific Oceans. Commun. Earth & Environ. 3, 121, DOI: 10.1038/s43247-022-00441-6 (2022).

11. Malard, L. A. et al. Aerobiology over the Southern Ocean – Implications for bacterial colonization of Antarctica. Environ. Int. 169, 107492, DOI: 10.1016/j.envint.2022.107492 (2022).

12. Wehking, J. et al. Community composition and seasonal changes of archaea in coarse and fine air particulate matter. Biogeosciences 15, 4205–4214, DOI: 10.5194/bg-15-4205-2018 (2018).

13. Šantl-Temkiv, T., Amato, P., Casamayor, E. O., Lee, P. K. H. & Pointing, S. B. Microbial ecology of the atmosphere. FEMS Microbiol. Rev. 46, fuac009, DOI: 10.1093/femsre/fuac009 (2022).

14. Cao, Y. et al. Airborne bacterial community diversity, source and function along the Antarctic Coast. Sci. Total. Environ. 765, 142700, DOI: 10.1016/j.scitotenv.2020.142700 (2021).

15. Alsante, A. N., Thornton, D. C. O. & Brooks, S. D. Ocean Aerobiology. Front. Microbiol. 12, 764178, DOI: 10.3389/fmicb.2021.764178 (2021).

16. Gusareva, E. S. et al. Microbial communities in the tropical air ecosystem follow a precise diel cycle. Proc. Natl. Acad. Sci. United States Am. 116, 23299–23308, DOI: 10.1073/pnas.1908493116 (2019).

17. Tignat-Perrier, R. et al. Global airborne microbial communities controlled by surrounding landscapes and wind conditions. Sci. Reports 9, 14441, DOI: 10.1038/s41598-019-51073-4 (2019).

18. Seinfeld, J. H. & Pandis, S. N. From air pollution to climate change. Atmospheric chemistry physics 1326 (1998).

19. Geever, M. Submicron sea spray fluxes. Geophys. Res. Lett. 32, DOI: 10.1029/2005GL023081 (2005).

20. Grythe, H., Ström, J., Krejci, R., Quinn, P. & Stohl, A. A review of sea-spray aerosol source functions using a large global set of sea salt aerosol concentration measurements. Atmospheric Chem. Phys. 14, 1277–1297, DOI: 10.5194/acp-14-1277-2014 (2014).

21. Griffin, D. W., Garrison, V. H., Herman, J. R. & Shinn, E. A. African desert dust in the Caribbean atmosphere: microbiology and public health. Aerobiologia 17, 203–213, DOI: 10.1023/A:1011868218901 (2001).

22. Wéry, N., Galès, A. & Brunet, Y. Microbiology of Aerosols, chap. 2.1, 115–135 (John Wiley Sons, Ltd, 2017). https://onlinelibrary.wiley.com/doi/pdf/10.1002/9781119132318.ch2a.

23. Prospero, J. M., Blades, E., Mathison, G. & Naidu, R. Interhemispheric transport of viable fungi and bacteria from Africa to the Caribbean with soil dust. Aerobiologia 21, 1–19, DOI: 10.1007/s10453-004-5872-7 (2005).

24. Šantl-Temkiv, T. et al. Characterization of airborne ice-nucleation-active bacteria and bacterial fragments. Atmospheric Environ. 109, 105–117, DOI: 10.1016/j.atmosenv.2015.02.060 (2015).

25. Wilson, T. W. et al. A marine biogenic source of atmospheric ice-nucleating particles. Nature 525, 234–238, DOI: 10.1038/nature14986 (2015).

26. Hamilton, D. S. et al. Earth, Wind, Fire, and Pollution: Aerosol Nutrient Sources and Impacts on Ocean Biogeochemistry. Annu. Rev. Mar. Sci. 14, 303–330, DOI: 10.1146/annurev-marine-031921-013612 (2022).

27. Mahowald, N. et al. Global distribution of atmospheric phosphorus sources, concentrations and deposition rates, and anthropogenic impacts. Glob. biogeochemical cycles 22, DOI: 10.1029/2008GB003240 (2008).

28. Polymenakou, P. N., Mandalakis, M., Stephanou, E. G. & Tselepides, A. Particle Size Distribution of Airborne Microorganisms and Pathogens during an Intense African Dust Event in the Eastern Mediterranean. Environ. Heal. Perspectives 116, 292–296, DOI: 10.1289/ehp.10684 (2008).

29. Šantl-Temkiv, T. et al. Bioaerosol field measurements: Challenges and perspectives in outdoor studies. Aerosol Sci. Technol. 54, 520–546, DOI: 10.1080/02786826.2019.1676395 (2020).

30. Lee, H. et al. Ipcc, 2023: Climate change 2023: Synthesis report, summary for policymakers. contribution of working groups i, ii and iii to the sixth assessment report of the intergovernmental panel on climate change. Technical Report 10.59327/IPCC/AR6-9789291691647.001 (2023). DOI: 10.59327/IPCC/AR6-9789291691647.001.

31. Lappan, R. et al. The atmosphere: a transport medium or an active microbial ecosystem? The ISME J. 18, DOI: 10.1093/ismejo/wrae092 (2024).

32. Schiebel, R. et al. Probing the open ocean with the research sailing yacht Eugen Seibold for climate geochemistry. J. Geophys. Res. Atmospheres DOI: 10.1029/2023JD040581 (2024).

33. Longhurst, A. R. Chapter 1 - toward an ecological geography of the sea. In Ecological Geography of the Sea, DOI: 10.1016/B978-012455521-1/50002-4 (Academic Press, 2007), 2nd edn.

34. Jensen, T. L., Kreidenweis, S. M., Kim, Y., Sievering, H. & Pszenny, A. Aerosol distributions in the North Atlantic marine boundary layer during Atlantic Stratocumulus Transition Experiment/Marine Aerosol and Gas Exchange. J. Geophys. Res. Atmospheres 101, 4455–4467, DOI: 10.1029/95JD00506 (1996).

35. Milke, F., Meyerjürgens, J. & Simon, M. Ecological mechanisms and current systems shape the modular structure of the global oceans’ prokaryotic seascape. Nat. Commun. 14, 6141, DOI: 10.1038/s41467-023-41909-z (2023).

36. Ruiz-Gil, T. et al. Airborne bacterial communities of outdoor environments and their associated influencing factors. Environ. Int. 145, 106156, DOI: 10.1016/j.envint.2020.106156 (2020).

37. Souza, F. F. et al. Influence of seasonality on the aerosol microbiome of the Amazon rainforest. Sci. The Total. Environ. 760, 144092, DOI: 10.1016/j.scitotenv.2020.144092 (2021).

38. Failor, K. C., Schmale, D. G., Vinatzer, B. A. & Monteil, C. L. Ice nucleation active bacteria in precipitation are genetically diverse and nucleate ice by employing different mechanisms. The ISME J. 11, 2740–2753, DOI: 10.1038/ismej.2017.124 (2017).

39. Hecker, M. & Völker, U. General stress response of Bacillus subtilis and other bacteria. In Advances in Microbial Physiology, vol. 44, 35–91, DOI: 10.1016/S0065-2911(01)44011-2 (2001).

40. Flemming, H.-C. & Wuertz, S. Bacteria and archaea on Earth and their abundance in biofilms. Nat. Rev. Microbiol. 17, 247–260, DOI: 10.1038/s41579-019-0158-9 (2019).

41. Flombaum, P. et al. Present and future global distributions of the marine Cyanobacteria Prochlorococcus and Synechococcus. Proc. Natl. Acad. Sci. 110, 9824–9829, DOI: 10.1073/pnas.1307701110 (2013).

42. Schattenhofer, M. et al. Latitudinal distribution of prokaryotic picoplankton populations in the Atlantic Ocean. Environ. Microbiol. 11, 2078–2093, DOI: 10.1111/j.1462-2920.2009.01929.x (2009).

43. Polovina, J. J., Howell, E. A. & Abecassis, M. Ocean’s least productive waters are expanding. Geophys. Res. Lett. 35, DOI: 10.1029/2007GL031745 (2008).

44. Zubkov, M. V., Sleigh, M. A., Burkill, P. H. & Leakey, R. J. Picoplankton community structure on the Atlantic Meridional Transect: a comparison between seasons. Prog. Oceanogr. 45, 369–386, DOI: 10.1016/S0079-6611(00)00008-2 (2000).

45. Barns, S. M., Takala, S. L. & Kuske, C. R. Wide Distribution and Diversity of Members of the Bacterial Kingdom Acidobacterium in the Environment. Appl. Environ. Microbiol. 65, 1731–1737, DOI: 10.1128/AEM.65.4.1731-1737.1999 (1999).

46. Zeng, Y. et al. Metagenomic evidence for the presence of phototrophic Gemmatimonadetes bacteria in diverse environments. Environ. Microbiol. Reports 8, 139–149, DOI: 10.1111/1758-2229.12363 (2016).

47. Slade, D. & Radman, M. Oxidative Stress Resistance in Deinococcus radiodurans. Microbiol. Mol. Biol. Rev. 75, 133–191, DOI: 10.1128/MMBR.00015-10 (2011).

48. Yilmaz, P., Yarza, P., Rapp, J. Z. & Glöckner, F.O. Expanding the World of Marine Bacterial and Archaeal Clades. Front. Microbiol. 6, 1524, DOI: 10.3389/fmicb.2015.01524 (2016).

49. Fuchs, B., Woebken, D., Zubkov, M., Burkill, P. & Amann, R. Molecular identification of picoplankton populations in contrasting waters of the Arabian Sea. Aquatic Microb. Ecol. 39, 145–157, DOI: 10.3354/ame039145 (2005).

50. Louca, S., Doebeli, M. & Parfrey, L. W. Correcting for 16S rRNA gene copy numbers in microbiome surveys remains an unsolved problem. Microbiome 6, 1–12, DOI: 10.1186/s40168-018-0420-9 (2018).

51. Fröhlich-Nowoisky, J. et al. Diversity and seasonal dynamics of airborne archaea. Biogeosciences 11, 6067–6079, DOI: 10.5194/bg-11-6067-2014 (2014).

52. Prass, M. et al. Bioaerosols in the Amazon rain forest: Temporal variations and vertical profiles of Eukarya, Bacteria and Archaea. Biogeosciences 18, 4873–4887, DOI: 10.5194/bg-2020-469 (2021).

53. Church, M. J. et al. Abundance and distribution of planktonic Archaea and Bacteria in the waters west of the Antarctic Peninsula. Limnol. Oceanogr. 48, 1893–1902, DOI: 10.4319/lo.2003.48.5.1893 (2003).

54. Willig, M., Kaufman, D. & Stevens, R. Latitudinal Gradients of Biodiversity: Pattern, Process, Scale, and Synthesis. Annu. Rev. Ecol. Evol. Syst. 34, 273–309, DOI: 10.1146/annurev.ecolsys.34.012103.144032 (2003).

55. Moss, J. A., Henriksson, N. L., Pakulski, J. D., Snyder, R. A. & Jeffrey, W. H. Oceanic Microplankton Do Not Adhere to the Latitudinal Diversity Gradient. Microb. Ecol. 79, 511–515, DOI: 10.1007/s00248-019-01413-8 (2020).

56. Thompson, L. R. et al. A communal catalogue reveals Earth’s multiscale microbial diversity. Nature 551, 457–463, DOI: 10.1038/nature24621 (2017).

57. Sunagawa, S. et al. Structure and function of the global ocean microbiome. Science 348, 1261359, DOI: 10.1126/science.1261359 (2015).

58. Fuhrman, J. A. et al. A latitudinal diversity gradient in planktonic marine bacteria. Proc. Natl. Acad. Sci. 105, 7774–7778, DOI: 10.1073/pnas.0803070105 (2008).

59. Aardema, H. M. et al. On the variability of phytoplankton photophysiology along a latitudinal transect in the North Atlantic surface ocean. J. Geophys. Res. Biogeosciences DOI: 10.1029/2023JG007962 (2024).

60. Guo, C. et al. Shifts in Microbial Community Structure and Activity in the Ultra-Oligotrophic Eastern Mediterranean Sea Driven by the Deposition of Saharan Dust and European Aerosols. Front. Mar. Sci. 3, 170, DOI: 10.3389/fmars.2016.00170 (2016).

61. Rahav, E., Belkin, N., Paytan, A. & Herut, B. The Relationship between Air-Mass Trajectories and the Abundance of Dust-Borne Prokaryotes at the SE Mediterranean Sea. Atmosphere 10, 280, DOI: 10.3390/atmos10050280 (2019).

62. Baker, A., Jickells, T., Biswas, K., Weston, K. & French, M. Nutrients in atmospheric aerosol particles along the Atlantic Meridional Transect. Deep. Sea Res. Part II: Top. Stud. Oceanogr. 53, 1706–1719, DOI: 10.1016/j.dsr2.2006.05.012 (2006).

63. Maki, T. et al. Aeolian Dispersal of Bacteria Associated With Desert Dust and Anthropogenic Particles Over Continental and Oceanic Surfaces. J. Geophys. Res. Atmospheres 124, 5579–5588, DOI: 10.1029/2018JD029597 (2019).

64. Gat, D., Mazar, Y., Cytryn, E. & Rudich, Y. Origin-Dependent Variations in the Atmospheric Microbiome Community in Eastern Mediterranean Dust Storms. Environ. Sci. & Technol. 51, 6709–6718, DOI: 10.1021/acs.est.7b00362 (2017).

65. Ruff, S. E. et al. A global comparison of surface and subsurface microbiomes reveals large-scale biodiversity gradients, and a marine-terrestrial divide. Sci. Adv. 10, 1–18, DOI: 10.1126/sciadv.adq0645 (2024).

66. Xu, A., Liu, C., Zhao, S., Song, Z. & Sun, H. Dynamic distribution of Massilia spp. in sewage, substrate, plant rhizosphere/phyllosphere and air of constructed wetland ecosystem. Front. Microbiol. 14, 1–17, DOI: 10.3389/fmicb.2023.1211649 (2023).

67. Dworkin, M., Falkow, S., Rosenberg, E.Schleifer, K.-H. & Stackebrandt, E. The Prokaryotes - A Handbook on the Biology of Bacteria (Springer, 2006).

68. Rahlff, J. et al. High wind speeds prevent formation of a distinct bacterioneuston community in the sea-surface microlayer. FEMS Microbiol. Ecol. 93, fix041, DOI: 10.1093/femsec/fix041 (2017).

69. O’Dowd, C. D. & Smith, M. H. Physicochemical properties of aerosols over the northeast Atlantic: Evidence for wind-speed-related submicron sea-salt aerosol production. J. Geophys. Res. Atmospheres 98, 1137–1149, DOI: 10.1029/92JD02302 (1993).

70. Wurl, O., Wurl, E., Miller, L., Johnson, K. & Vagle, S. Formation and global distribution of sea-surface microlayers. Biogeosciences 8, 121–135, DOI: 10.5194/bg-8-121-2011 (2011).

71. Jang, J. et al. Selective transmission of airborne bacterial communities from the ocean to the atmosphere over the Northern Pacific Ocean. Sci. The Total. Environ. 957, 177462, DOI: 10.1016/j.scitotenv.2024.177462 (2024).

72. Salazar, G. et al. Gene Expression Changes and Community Turnover Differentially Shape the Global Ocean Metatranscriptome. Cell 179, 1068–1083, DOI: 10.1016/j.cell.2019.10.014 (2019).

73. Vuillemin, A., Coskun, Ö.K. & Orsi, W. D. Microbial Activities and Selection from Surface Ocean to Subseafloor on the Namibian Continental Shelf. Appl. Environ. Microbiol. 88, e00216–22, DOI: 10.1128/aem.00216-22 (2022).

74. Li, L. et al. Bacterial communities in cropland soils: Taxonomy and functions. Plant Soil 1–19, DOI: 10.1007/s11104-023-06396-7 (2023).

75. DeBruyn, J. M., Nixon, L. T., Fawaz, M. N., Johnson, A. M. & Radosevich, M. Global Biogeography and Quantitative Seasonal Dynamics of Gemmatimonadetes in Soil. Appl. Environ. Microbiol. 77, 6295–6300, DOI: 10.1128/AEM.05005-11 (2011).

76. Michaud, J. M. et al. Taxon-specific aerosolization of bacteria and viruses in an experimental ocean-atmosphere mesocosm. Nat. Commun. 9, 2017, DOI: 10.1038/s41467-018-04409-z (2018).

77. Hu, D., Cha, G. & Gao, B. A Phylogenomic and Molecular Markers Based Analysis of the Class Acidimicrobiia. Front. Microbiol. 9, 987, DOI: 10.3389/fmicb.2018.00987 (2018).

78. Dadon-Pilosof, A. et al. Surface properties of SAR11 bacteria facilitate grazing avoidance. Nat. Microbiol. 2, 1608–1615, DOI: 10.1038/s41564-017-0030-5 (2017).

79. Morris, R. M. et al. SAR11 clade dominates ocean surface bacterioplankton communities. Nature 420, 806–810, DOI: 10.1038/nature01240 (2002).

80. Giovannoni, S. J. SAR11 Bacteria: The Most Abundant Plankton in the Oceans. Annu. Rev. Mar. Sci. 9, 231–255, DOI: 10.1146/annurev-marine-010814-015934 (2017).

81. Brüwer, J. D. et al. In situ cell division and mortality rates of SAR11, SAR86, Bacteroidetes, and Aurantivirga during phytoplankton blooms reveal differences in population controls. mSystems e01287-22, DOI: 10.1128/msystems.01287-22 (2023).

82. Freitas, G. P. et al. Emission of primary bioaerosol particles from Baltic seawater. Environ. Sci. Atmospheres 2, 1170–1182, DOI: 10.1039/D2EA00047D (2022).

83. Aller, J. Y., Kuznetsova, M. R., Jahns, C. J. & Kemp, P. F. The sea surface microlayer as a source of viral and bacterial enrichment in marine aerosols. J. Aerosol Sci. 36, 801–812, DOI: 10.1016/j.jaerosci.2004.10.012 (2005).

84. von der Weiden, S.-L., Drewnick, F. & Borrmann, S. Particle Loss Calculator – a new software tool for the assessment of the performance of aerosol inlet systems. Atmospheric Meas. Tech. 2, 479–494, DOI: 10.5194/amt-2-479-2009 (2009).

85. Slagter, H. A. et al. Semicontinuous phytoplankton flowcytometry from the Norwegian Sea into the equatorial upwelling., DOI: 10.17617/3.MLNQJ4 (2023).

86. Stein, A. F. et al. NOAA’s HYSPLIT Atmospheric Transport and Dispersion Modeling System. Bull. Am. Meteorol. Soc. 96, 2059–2077, DOI: 10.1175/BAMS-D-14-00110.1 (2015).

87. Draxler, R. R. & Hess, G. D. An overview of the HYSPLIT_4 modelling system for trajectories, dispersion and deposition. Aust. Meteorol. Mag. 47, 295–308 (1998).

88. Caporaso, J. G. et al. Ultra-high-throughput microbial community analysis on the Illumina HiSeq and MiSeq platforms. ISME J. 6, 1621–1624, DOI: 10.1038/ismej.2012.8 (2012).

89. Apprill, A., McNally, S., Parsons, R. & Weber, L. Minor revision to V4 region SSU rRNA 806R gene primer greatly increases detection of SAR11 bacterioplankton. Aquatic Microb. Ecol. 75, 129–137, DOI: 10.3354/ame01753 (2015).

90. Wang, F. et al. Assessment of 16S rRNA gene primers for studying bacterial community structure and function of aging flue-cured tobaccos. AMB Express 8, 182, DOI: 10.1186/s13568-018-0713-1 (2018).

91. Klindworth, A. et al. Evaluation of general 16S ribosomal RNA gene PCR primers for classical and next-generation sequencing-based diversity studies. Nucleic Acids Res. 41, e1–e1 (2013).

92. Bennke, C. M. et al. Modification of a High-Throughput Automatic Microbial Cell Enumeration System for Shipboard Analyses. Appl. Environ. Microbiol. 82, 3289–3296, DOI: 10.1128/AEM.03931-15 (2016).

93. R Core Team. R: A Language and Environment for Statistical Computing. R Foundation for Statistical Computing (2023).

94. Callahan, B. J. et al. Dada2: High-resolution sample inference from illumina amplicon data. Nat. Methods 13, 581–583, DOI: 10.1038/nmeth.3869 (2016).

95. Davis, N. M., Proctor, D., Holmes, S. P., Relman, D. A. & Callahan, B. J. Simple statistical identification and removal of contaminant sequences in marker-gene and metagenomics data. Microbiome 6, 1–14 (2018).

96. Heintzenberg, J., Covert, D. C. & Van Dingenen, R. Size distribution and chemical composition of marine aerosols: a compilation and review. Tellus B: Chem. Phys. Meteorol. 52, 1104–1122, DOI: 10.3402/tellusb.v52i4.17090 (2000).

97. Dedrick, J. L., Saliba, G., Williams, A. S., Russell, L. M. & Lubin, D. Retrieval of the sea spray aerosol mode from submicron particle size distributions and supermicron scattering during LASIC. Atmospheric Meas. Tech. 15, 4171–4194, DOI: 10.5194/amt-15-4171-2022 (2022).

