## Supplementary Materials for "Latitudinal diversity gradients and selective microbial exchange at the Atlantic ocean–air interface"

### Supporting Results and Discussion

##### *Detailed description of community composition of water and air communities*

*Flavobacteriales.* Flavobacteriales in the phylum Bacteroidota (Bacteroidetes) and the class Bacteroidia, are suggested to relate to a higher primary productivity, correlating with phytoplankton blooms (1). As highlighted in the main text, Flavobacteriales were the dominant order level clade in the water and also present in all air samples. Their significantly higher relative sequence abundance (RSA) in the mesotrophic northern regions compared to the oligotrophic southern regions in both, air and water communities supports the previously found association with both, colder (2) and nutrient-rich waters exhibiting a higher primary productivity. However, in the water samples, after dropping to a minimal RSA of 20 % between 38° N – 29° N, RSA increased again and stabilized at around 36 % between 21° N – 3° N. The air community showed a similar decrease in RSA of Flavobacteriales, from 67° N to 38° N, however, remained low afterwards. This might be due the very high RSAs of Firmicutes (Bacillota) in the air in this area.

*Additional phyla significantly associated with the air microbiome* Planctomycetota, Gemmatimonadota and Deinococcota were all significantly higher abundant in the air ($p<0.05$, $p<0.001$, $p<0.001$, respectively).

*Additional phyla significantly associated with the water microbiome* Marinimicrobia ($p<0.001$), Verrucomicrobiota ($p<0.001$), the archaeal phylum Thermoplasmatota, Bdellovibrionota ($p<0.001$), Dadabacteria (Dadabacteriota, $p<0.001$), Desulfobacterota ($p<0.001$), SAR324 ($p<0.001$) and Margulisbacteria (Margulisiibacteriota, $p<0.001$, Fig. S4).

*Archaeal composition of water and air communities.* Archaeal biodiversity and relative importance in the ocean is suggested to increase with depth (3). Also, a negative correlation of dissolved oxygen and chlorophyll-*a* fluorescence with the abundance of Archaea within the water column has been described (4). All surface water communities within the sampled transect showed archaeal sequence abundance, however, they were only present in 13 of 36 air samples (39 % prevalence). We are aware of the limitations set by the usage of a universal primer pair to target Bacteria and Archaea simultaneously, however, intriguingly, the air and water community showed highly significant differences in the relative abundance of Archaea (Wilcoxon $p<0.0001$, MaAsLin2 coef. 6.8, q-value<0.0001). In addition, Archaeal RSA increased towards the equator, in contrast to what was previously described (5).

### Supporting Methods

#### Microbial air sampling – Detailed sampling protocol

All sampling equipment underwent thorough cleaning inside a biological safety cabinet under laminar flow (MSC-Advantage$^{TM}$ Class II Biological Safety Cabinet, Thermo Fisher Scientific$^{TM}$) using a 70 % ethanol solution. All laboratory surfaces used for sample processing were wiped with 70 % ethanol right before and during sample processing. At all times, either nitrile or latex gloves were worn and repeatedly cleaned with 70 % ethanol during sample processing. To monitor and account for any remaining contamination, sampling blanks were regularly taken and processed. All parts of the air sampler (Coriolis µ) in direct contact with the sample were cleaned using 70 % ethanol before and after sampling and rinsed with ultra-pure water (MQ; resistivity 18.2 MΩ·cm) prior to each sample. While not in use, all parts of the Coriolis µ potentially contributing to contamination were rinsed with 70 % ethanol and stored in clean zip-log bags. Blanks were taken using the same collection liquid but without air collection. Liquid was supplied through the continuous liquid supply system to also check for possible contamination introduced by the continuous liquid supply system of the Coriolis µ. Samples and Blanks were collected using 1×phosphate buffered saline (PBS) with MQ as continuous liquid supply or in MQ directly, both filtered through a 0.1 µm pore size filter. Liquid losses due to evaporation were compensated by adding MQ with a flow-rate between 0.1 and 0.7 mL per min, depending on meteorological conditions. In addition to the described samples, an additional sample for ice nucleation assays/cell culture was taken from the air samples. For those, 0.5 to 1 mL of the sample were either mixed with an equal volume of Glycerol (f.c. 25 % Glycerol, 1xPBS) or left untreated and then frozen at -80 °C. For water samples, 1 to 2 mL of the sample were either mixed with an equal volume of Glycerol (f.c. 25 % Glycerol, 1xPBS) or left untreated and then frozen at -80 °C for cell culture/ice nucleation assays. These samples are not part of this study.

#### DNA extraction – Kit comparison and modifications

A summary of the results of the tested protocols can be found in SI Appendix, Dataset S7. In short, 15 samples and 4 blanks were collected as described above, microbial cell abundance and particle load of filters was investigated using DAPI-stained filters (cf. Materials and Methods, Staining of microbial cells). Filters were grouped in "clean", "medium" and "dirty", depending on their cell and particle load. For each condition, filters were cut in 4 pieces and distributed amongst the 4 protocols to be tested, resulting in one whole filter per assay (consisting of 4 filter pieces, 1 from each filter belonging to the respective group).

**Table 1.** Relative Sequence Abundance (RSA) on phylum level for top 11 phyla ordered by decreasing mean RSA (in%) and selected lineages in overall sample set (air and water samples). Rank: Position in each data set separately when ordered by decreasing mean RSA. Value for "Other" in the Rank column represents the number of remaining phylum level clades in the respective biome.

| Phylum | Rank Air | RSA Air | Rank Water | RSA Water |
|---|---|---|---|---|
| Proteobacteria (Pseudomonadota) | 1 | 37.6 | 1 | 39 |
| Bacteroidota (Bacteroidetes) | 3 | 17.8 | 2 | 34.9 |
| Cyanobacteria | 4 | 11.8 | 3 | 14.8 |
| *Synechococcus* | | 8.4 | | 7.7 |
| *Prochlorococcus MIT9313* | | 1 | | 6.4 |
| Firmicutes (Bacillota) | 2 | 24.3 | 21 | 0.005 |
| Actinobacteriota (Actinomycetota) | 5 | 3.2 | 5 | 2.2 |
| Marinimicrobia-SAR406- | 16 | 0.1 | 4 | 2.5 |
| Planctomycetota | 6 | 1.5 | 8 | 1.0 |
| Verrucomicrobiota | 14 | 0.2 | 6 | 1.5 |
| Thermoplasmatota (Arc) | 20 | 0.04 | 7 | 1.2 |
| Bdellovibrionota | 10 | 0.3 | 9 | 0.7 |
| Acidobacteriota | 7 | 1 | NA | NA |
| Other | n=30 | 2.1 | n=13 | 2.2 |
| Air remaining top 10 | | | | |
| Deinococcota | 8 | 0.5 | NA | NA |
| Gemmatimonadota | 9 | 0.4 | NA | NA |
| Water remaining top 10 | | | | |
| Dadabacteria | 18 | 0.05 | 10 | 0.6 |

Only for the "dirty" filters, three filters were used, resulting in 3 quarter filter pieces per assay. DNA was extracted using commercially available DNA extraction kits (1. ChargeSwitch™ Forensic DNA Purification Kit (ThermoFisher Scientific), 2. DNeasy® PowerSoil® Pro Kit (Qiagen), 3. Quick-DNA/RNA™ Microprep Plus Kit (Zymo Research)) and 4. a modified chloroform extraction protocol after (7). The ChargeSwitch™ Forensic DNA Purification Kit (ThermoFisher Scientific) was found to be most efficient and samples presented in this study were extracted with the following modifications. Air filters were inserted in a 2 mL centrifuge tube as a whole with the sampled side facing to the inside of the tube. Water filters with a diameter of 25 mm were cut in half, and half of the filter inserted in a 2 mL centrifuge tube with the sampled side facing the inside of the tube. Water filters with a diameter of 47 mm were cut in pieces and the pieces inserted in a 2 mL centrifuge tube. The following steps were performed for all samples. After adding 1 mL of lysis buffer the sample-lysis buffer mixture was thrice frozen at -20 °C and thawed. Afterwards, 50 µL of lysozyme (stock concentration 50 mg $mL^{-1}$) were added, the mixture inverted thrice, and incubated for 1 h at 37 °C, 450 rpm with a short vortex after 30 min. Afterwards, 15 µL Proteinase K (20 mg $mL^{-1}$ in 50 mM Tris-HCl, pH 8.5, 5 mM $CaCl_2$, 50 % glycerol) were added and the mixture incubated for 90 min at 55 °C, 450 rpm with a short vortex after 45 min. Afterwards, samples were vortexed for 5-15 sec, inverted thrice, and shortly spinned down before all liquid was transferred to a fresh 2 mL centrifuge tube. Binding and washing steps were performed as described in the manufacturers protocol. For the elution step, 100 µL of elution buffer was added and incubated at room temperature for 5 min (shortly pipetted up and down after 2.5 min). The eluate was transferred to a clean tube (final DNA product) and the elution process repeated, resulting in a final volume of DNA eluate of 200 µL. DNA concentration was determined using the DeNovix dsDNA Ultra High Sensitivity quantification assay (DeNovix) and measured on a Qubit™ fluorometer. A strong correlation (Pearson correlation) between DNA yield and cell numbers throughout sample classes (low-biomass air samples to high-biomass water samples) underline the suitability of this extraction method for the given sample set. For air samples, the volume of the final eluate was reduced from 200 µL to approx. 40 µL by incubation at 60 °C, 300 rpm to increase DNA concentration for subsequent sequencing. DNA eluate was stored at -70 (https://www.freezerchallenge.org/). All DNA extraction steps were performed in a clean environment with protective gloves under laminar flow (MSC-Advantage™ Class II Biological Safety Cabinet, Thermo Fisher Scientific™). All surfaces and gloves were cleaned with 70 % isopropanol, materials were autoclaved and cleaned with 70 % isopropanol before usage. A distinguished set of pipettes together with sterile filter-tip pipette tips were used to minimize the potential of sample contamination during DNA extraction.

**Table 2.** Sampled latitudinal provinces, after Longhurst, 2007 (6)

| Province Abbreviation | Province Name | Biome | Trophy | Latitude max ° N | Latitude min ° N |
|---|---|---|---|---|---|
| ARCT | Atlantic Arctic | polar | mesotrophic | 67 | 65 |
| SARC | Atlantic Sub-Arctic | polar | mesotrophic | 65 | 58.5 |
| NADR | North Atlantic Drift | westerly | mesotrophic | 58.5 | 43.5 |
| NASE | North Atlantic Subtropical Gyre | westerly | oligotrophic | 43.5 | 25 |
| NATR | North Atlantic Tropical Gyral | trade wind | oligotrophic | 25 | 12 |
| WTRA | Western Tropical Atlantic | trade wind | oligotrophic | 12 | 3 |

### Modified chloroform extraction protocol

For the modified chloroform extraction (7), 4 mL of Extraction Buffer (100 mM Tris-HCl, pH 8.0, 100 mM EDTA, pH 8.0, 100 mM sodium phosphate buffer, pH 8.0, 1.5 M NaCl, 1 % CTAB (cetyltrimethylammonium bromide)) were added to the filter and the sample vortexed, samples were frozen at -20 °C for 1 h or until they were frozen completely and then thawed at 65 °C. The freeze-thaw cycle was repeated thrice. After cooling down to 37 °C, 50 µL of 20 mg $mL^{-1}$ Proteinase K were added and the sample incubated overnight at 37 °C under mild horizontal shaking (200 rpm, 45° angle). The next day, 0.5 mL 20 % SDS (sodium dodecyl sulfate) were added, and the sample incubated at 65 °C for 60 min, gently inverting end over end every 20 min. The sample was centrifuged for 10 min, 4649×g at RT and the supernatant transferred to a new tube. An equal volume of Isopropanol/Chloroform (24:1) was added, the tube mixed thoroughly, the sample centrifuged at 4649×g at RT and the upper, aqueous phase collected. 0.6 volumes of isopropanol (abs.) were added and the eluate left at 4 °C over night to precipitate the DNA. As precipitate was visible (likely from CTAB), samples were heated to 65 °C shortly before centrifuging at 4649×g at RT for 1 h. The supernatant was discarded and the invisible pellet washed with 0.5 mL cold ethanol (80 %, v/v). The pellet was centrifuged at 4649×g at RT for 1 h, the supernatant decanted and the pellet dried for 20 min at RT in a clean environment. Finally, the pellet was re-suspended in 50 µL PCR-grade water and incubated for 3 h for the DNA to elute before being stored at -20 °C until concentration determination.

### Bioinformatic procedures

All analyses and visualizations were performed in the software environment R (8), using several packages (9; 10; 11; 12; 13; 14; 15; 16; 17; 18; 19; 20; 21; 22; 23; 24; 25; 26; 27; 28; 29; 30; 31; 32; 33; 34), together with custom scripts and modifications (https://github.com/bellahrabe/Microbial-community-analysis-Eugen2021).

#### *Primer properties*

Allowing for 1 mismatch with a 5 bases 0-mismatch zone at the 3' end, the used 16S v4-v5 primer pair covered 90.9 % of bacterial, 89.2 % of archaeal and 1.7 % of eukaryotic sequences of the SILVA RefNR SSU r138.1 database (0 mismatches: 81.4 % of all bacterial, 81.1 % of all archaeal and 0.1 % of all eukaryotes) at the time of analysis, as determined using TestPrime 1.0. A full list of lineage coverage can be found in Dataset S8.

#### *Illumina read processing and taxonomy assignment*

Initial read processing as well as the testing of contamination removal strategies was done with all samples of the sequencing run (including samples not part of this study). For 16S rRNA gene analysis, the MiSeq platform is known to miscalculate quality values for low-diversity libraries due to the miscalculation of the phasing/prephasing, which in turn leads to (unreasonably) bad Q30 values. Hence, quality trimming was not advised for the 16S rRNA reads. To ensure high-quality reads as well as reasonable ratios of reads passing the DADA2 pipeline, different trimming and error rate scenarios were tried (Dataset S9). Finally, reads were trimmed to 260 (forward) and 190 (reverse) bases. 19 and 20 bases were trimmed from the start of the forward and reverse reads respectively, to remove primer sequences. maxEE was set to c(3,6). Applying these filtering parameters resulted in 74 % of the reads passing through the DADA2 pipeline (Dataset S9). Taxonomy was assigned using the DADA2 algorithm with the SILVA138.1 data base (35; 36).

#### *Bioinformatic contamination removal strategies*

Potential contaminant sequences were filtered using different strategies and the effect of each strategy was investigated. Each final decontamination strategy (Combinations 1-5, Table 3) does consist of 4 or 5 succeeding steps, which themselves, can be parameterised. The result of each step by itself on the original data set was also investigated, in order to determine unwanted effects on the data set. The summarised results of each decontamination step as well as of Combinations 1-5 are listed in

Dataset S10.
For the first and second step, sample-blank groupings were assigned depending on the respective instrument/collection method (st; air sample blanks to air samples, water sample blanks to water samples), the respective instrument/collection method per year (stay) or without batches (all), assigning all blanks to all samples.
The first step consisted of the usage of the R package decontam (30). As our air samples are considered low biomass, we tested the efficiency of the package using both, the "normal" biomass (decontam::isContaminant) and the low-biomass approach (decontam::isNotContaminant). Sample-blank groupings were assigned using either stay or st blank grouping.

**Table 3.** Tested combinations of decontamination strategies. For all strategies, potential human contaminants and reagent contaminants (if showing an inverse correlation to the DNA concentration) were removed.

| Combination | sample-blank batch | decontam | 2nd step | remove additional pot. contaminants | ASVs passing through | Reads passing through |
|---|---|---|---|---|---|---|
| 1 | stay | high biomass | remove | F | 75 % | 43 % |
| 2 | st | high biomass | remove | F | 68 % | 37 % |
| 3 | st | low biomass | remove | F | 59 % | 33 % |
| 4 | st | low biomass | smax | F | 64 % | 49 % |
| 5 | st | low biomass | remove | T | 52 % | 28 % |

In the second step, ASVs found in blanks were either removed (r), or their maximum read count subtracted (smax) from the samples, with reads normalized for equal sequencing depth per sample (dividing each ASV read count by the sum of the read count per sample, so that the sum of all reads per sample equals 1). For this step the same sample-blank groupings were assigned as in step 1. In the third step we removed all potential human contaminants as found in several previous studies (Dataset S11), followed by the fourth step, where we investigated known reagent contaminants as described in (37) to exclude clades which have a strong negative correlation to the measured DNA concentration. As all samples were processed following the same protocol and using the same reagents, only trends occurring within the whole data set were considered, which resulted in no exclusion of potential reagent contaminants (Dataset S13). For Combination 5, we added an additional step, removing all ASVs belonging to a very exhaustive list of potential reagent contaminants (Dataset S11).

#### *Considerations for bioinformatic contamination removal strategies*

We consider each decontamination step and strategy has its strengths, however, based on our observations of the effects the different parameters had on our sample set, we consider Combination 4 (Table 3) to be most efficient in removing contaminating sequences, while impacting community structure least, for the following reasons.

*Blank-sample batch grouping:* We decided to process the samples using sample-blank batches where all blanks taken with one collection method (i.e., water and air samples separately) in order to have enough blanks per sample group (air:9, water:6) and to not introduce misleading trends by correcting with blanks from different sampling methodologies.

*decontam package:* After removing potential contaminants using the decontam package with varying parameters, a lot of different ASVs and reads were still present in the blank samples (19–89 % of ASVs and 12–81 % of reads) with the least proportions passing through when using the low biomass approach. Hence, we consider the usage of the package a valuable additional step, but see the need of additional steps to efficiently remove contaminant sequences.

*Normalization:* Normalization of reads for equal sequencing depth did not have a substantial effect on the proportion of ASVs or reads passing through the decontamination process in all cases. Depending on the remaining filtering parameters, both, an increase or decrease in reads passing through, could be observed. As we consider a normalization step beneficial for reproducibility of results and applicability to different data sets, samples and reads were normalized (the decontam package is normalizing reads internally, hence, a normalization was only necessary for the manual decontamination step 2).

*Statistical test:* Applying a statistical test (here: Wilcoxon ranked-sum test) to identify contaminant samples led to overall more left-over ASVs and reads in the blank samples and was hence did not considere it appropriate for this data set. This might be due to several limitations of the data set, relevant for the success of this kind of statistical test, such as the limited sample size. Also, while the wilcoxon test does not assume a normal distribution, a difference in the compared distributions in the blanks and samples (which can be expected) might impact the outcome. In particular, when removing all reads occurring in any of the ASVs present in the blanks, the proportion of ASVs and reads in the samples after removal is much lower when defining contaminant ASVs with the wilcoxon test, supporting the assumption that results from the test are flawed for this kind of data set.

*ASV removal vs subtraction of their maximum read count vs subtraction of their average read count:* While subtracting the average read count observed in the blanks from the samples leads to a high proportion of reads and ASVs kept in the samples, also ASVs and reads kept in the blanks is high. This suggests that this approach is not suitable to catch most of the contaminant ASVs and reads. Simply removing all ASVs present in the blanks leads to the highest loss of ASVs and reads (11–82 % of ASVs and 51–71 % of reads, depending on the other filtering parameters). It might lead to the removal of clades which are present in the samples at high abundances and occur in one of the blanks at a very low abundance due to carry-over in any of the processing steps, and might therefore introduce trends not representative for the collected biome. Hence, we consider subtracting the maximum occurrence of an ASV in any of the blanks (belonging to the respective sampling methodology) a good compromise between loosing true diversity and trends and efficient removal of potential contaminant sequences.

***Results of bioinformatic contamination removal combinations 1-5***

Combinations 1 and 2 are similar to the decontamination strategy used for analysing a low-biomass air data set before (37). For Combination 1, we grouped blanks and samples separately by sampling technique and by sampling year, in Combination 2, we only grouped them by sampling technique. This resulted in the additional loss of only 7 % of our ASVs, and 6 % of of our reads (Table 4), indicating that the blank communities of different years showed rather similar ASVs. As sampling took place within 2 subsequent years, using the same setup and sampling materials, we argue that using a broader blank grouping (by technique, not by year) will result in a better identifications of potential contaminant sequences and hence continued with this blank-sample grouping for the following combinations.

**Table 4.** ASVs and reads (ASVs/reads, in %) passing through the different decontamination steps in the five decontamination strategies

| | 1 | 2 | 3 | 4 | 5 |
|---|---|---|---|---|---|
| 1) decontam | 92 / 85 | 93 / 85 | 81 / 76 | 81 / 76 | 81 / 76 |
| 2) removal/subtraction | 76 / 44 | 71 / 39 | 61 / 35 | 66 / 52 | 61 / 35 |
| 3) human contaminants | 75 / 43 | 68 / 18 | 59 / 33 | 64 / 49 | 59 / 33 |
| 4) reagent contaminants | 75 / 43 | 68 / 18 | 59 / 33 | 64 / 49 | 59 / 33 |
| 5) contaminants extended | | | | | 52 / 28 |

For Combinations 3–5 we used the low-biomass approach of the decontam package, followed by the removal (combination 3&5) or the subtraction of the maximum read occurrence of the normalized reads (combination 4) occurring in the blanks. Compared to the high-biomass approach used in combinations 1&2 we observed a slightly higher loss of ASVs and reads in our samples (Table 4) together with a significantly higher loss of ASVs and reads in our blanks (Table 5). This was especially pronounced for the blanks of our low-biomass samples. As an efficient decontamination of the low-biomass samples is of utmost importance, we suggest that using the low-biomass approach of the decontam package is more appropriate for our data set. Combination 3 and 4 solely differ in the second step, where combination 3 removes all ASVs found in the respective blank groups, while combination 4 subtracts the maximum value of reads in each ASV found in the respective blank groups from the samples. By subtracting the maximum read count and not removing those ASVs, we account for the potential carry-over of small sample volumes and cross-contamination during sample-processing. We believe that this approach better reflects true trends observable in the sampled ecosystem and eliminates the risk to entirely loose clades due to minor (cross-)contamination. Combination 5 serves as an additional control. Here, in addition to step 4 (removal of potential reagent contaminants in case they show an inverse correlation with the DNA concentration) we removed all ASVs/reads belonging to clades previously found as potential contaminants, in particular in studies dealing with low-biomass samples (38; 39, Dataset S11). We are aware that the result of this combination unlikely represents a real-life scenario, however, results of this combination were used to investigate whether trends seen within this study also held true if this very stringent decontamination strategy was applied.

**Table 5.** ASVs and reads (in %) passing through the first decontamination step (decontam) in the blank samples, using the normal (1 and 2) and low-biomass strategy (3-5) and different sample blank groupings.

| | 1 | 2 | 3-5 |
|---|---|---|---|
| 1) decontam | 71 / 56 | 74 / 66 | 31 / 25 |

Based on the above argumentation we propose combination 4 to be efficient in decontaminating the presented sample set. This approach proved to be efficient in detecting and removing ASVs and reads which were found in particular in our low-biomass blanks. In addition, potential human contaminants were mostly identified in the first two steps already, resulting in only a minor additional loss of ASVs/reads when manually removing previously known classical human contaminants. In addition, subtracting the normalized read occurrence instead of removing ASVs completely promises to leave true, natural community trends unaltered. However, we are aware of the risk of introducing misleading trends with the applied filtering methods. Therefore, we consistently verified our results by checking both, the original data set (no filtering), and the most stringent (combination 5) filtering approach. With this validation process we ensure the reliability of our conclusions, adding confidence to our findings, mitigating potential biases and errors that may (or may not) arise from the applied filtering methods.

#### *Classification of microbial lineages as air or ocean associated*

Microbial lineages were classified as either air associated or ocean associated or similarly associated with both biomes. Also, the potential impact of these organisms on the other biome was assigned, resulting in a total of 8 groups. The applied rule-set is summarised in Tbl. 6. The RSA was considered to be higher in one of the biomes (lower atmosphere (air), surface ocean (ocean)) when the $p$-value of the paired Wilcoxon signed rank test was lower than 0.05.

**Table 6.** Rule-set applied to classify microbial lineages to one of the groups.

| Group | Description | RSA | Correlation | RSA Air | RSA Sea | Prevalence Air | Prevalence Sea |
|---|---|---|---|---|---|---|---|
| Air to Ocean | Air associated, impacting surface ocean communities | Air>Sea | + | | | | |
| Air | Air associated, not impacting surface ocean communities | Air>Sea | not + | | <0.5 % | | <0.3 |
| AqO | Air associated, possibly impacting surface ocean communities | Air>Sea | not + | | >0.5 % | | >0.3 |
| Ocean to Air | Surface ocean associated, impacting air communities | Air<Sea | + | | | | |
| Ocean | Surface ocean associated, not impacting air | Air<Sea | not + | <0.5 % | | <0.3 | |
| OqA | Surface ocean associated, possibly impacting air communities | Air<Sea | not + | >0.5 % | | >0.3 | |
| Both | Associated with both biomes, likely exchanged | Air=Sea | + | | | | |
| BBq | Associated with both biomes, likely not exchanged | Air=Sea | not + | | | | |

### Cell enumeration

#### *Automated image recording – Modifications*

Images were recorded as described in (40) with some modifications accounting for the complexity of the air samples. For all samples, pictures using the 385 nm LED (DAPI) and the 590 nm LED (autofluorescence) were used. For air samples pictures using the 469 nm LED (potential autofluorescence of dust particles) were taken in addition (explanation below). Depending on the sample, 105 to 190 fields of view (FOVs) per sample were identified with a 1×magnification.

#### *Image processing and analysis using the ACME tool – Modifications*

*Water samples* For water samples (DAPI and autofluorescence channel), the following settings were used for automated cell detection. DAPI-channel set definition: Area>16 and Area<200 and SBR>2, DAPI-channel sub-set definition: NR_auto500ms_Signals=0, autofluorescence-channel (auto500ms) set definition: Area>16, DAPI to auto500ms overlap definition: Percent>40. Pictures were screened and detection of cells was considered good for all samples and no signals were added manually.

*Air samples* Due to the diverse nature and complexity of the air samples, these definitions were not considered suitable for the sample set after manual inspection of detected/not detected signals. To increase data quality, one additional channel (fish50ms/fish40ms: excitation using the 469 nm LED) was recorded, to exclude signals showing an autofluorescence when excited with this source. In addition, for a subset of samples the DAPI signal was recorded twice (DAPI and DAPI_auto) as using an automated excitation time seemed to result in higher quality pictures. The following settings were found to be most suitable for the majority of the samples and were used throughout the sample set to increase consistency within. DAPI-channel set definition: Area>12 and Area <100 and SBR>1.7, DAPI-channel sub-set definition: Nr_auto500ms_Signals=0 and Nr_fish40ms_Signals=0. DAPI_auto-channel (if present) set definition: Area>16 and Area <200 and SBR>2 and subset definition: Nr_auto500ms_Signals=0. These conditions were set to use this channel as a direct comparison to the water sample set (same set and subset parameters), while the other DAPI channel was adapted in order to catch more signals which were also considered cells by the human eye. The autofluorescence (auto500ms, fish40ms, fish50ms, fish60ms) set definitions were set to Area>12 throughout. The overlap definition between both DAPI channels and all autofluorescence channels were set to Percent>40.

#### *Cell number calculation*

The sample report returns the counted signals per number of FOVs, whereas the FOV report returns the number of counted signals for each counted FOV separately. One FOV counting frame has a width of 1348 and a height of 1000 pixels. Here, one pixel equals 0.1016 µm in the filter sample, hence, 1 counting frame covers a filter sample area of ~ 0.0139 $mm^2$. The proportion of the respective counting frames were calculated and the average cell number per grid divided by this proportion in order to derive the cell number per filter. For air samples, this number was then divided by the filtered air volume to derive cells per $m^3$. Usually, 1.5 mL of a 15 mL sample were filtered on a 25 mm filter (filter radius: r = 10 mm). These 15 mL samples were usually collected over 6 h, with a pump speed of 300 lpm, hence, 1.5 mL of sample equal 10.8 $m^3$ of collected air. For water samples, the derived cell number was divided by the filtered volume of collected sea water (usually 19 mL) to derive cells per mL.

**Table 7.** Summary of cell abundances in the lower atmosphere (~3.5 above the ocean surface) over the North East Atlantic Ocean in cells per $m^3$ for corrected and uncorrected cell abundances. These values can be considered lower and upper limits. And cells per mL observed in the surface ocean (~3 water depth).

| | cells $m^{-3}$ | cells $m^{-3}$ corrected | cells $10^6$ $mL^{-1}$ |
|---|---|---|---|
| Average | $1.2 \times 10^5$ | $8.2 \times 10^5$ | 1 |
| Standard Deviation | $2.1 \times 10^5$ | $1.5 \times 10^6$ | 0.4 |
| Minimum | 0 | 0 | 0.4 |
| 1.Quartile | $5.7 \times 10^1$ | $2.6 \times 10^2$ | 0.6 |
| Median | $5.0 \times 10^2$ | $2.3 \times 10^3$ | 0.9 |
| 3.Quartile | $1.6 \times 10^5$ | $1.2 \times 10^6$ | 1.1 |
| Maximum | $7.4 \times 10^5$ | $4.8 \times 10^6$ | 2.3 |

#### *Correction of microbial abundances in low-biomass samples*

Due to the very low cell numbers in the atmospheric samples and therefore on the stained filters, very few FOVs were recorded for several samples (true for several air samples and water and air blanks). This is considered to be due to a lack of signal and not due to other technical limitations. After visual inspection of the recorded pictures, samples with very few recorded FOVs due to the above reasons were corrected as follows. The average ratio between recorded pictures and set pictures to be taken for high-quality samples was calculated (0.95 ± 0.15, mean ± standard deviation). This average was used to calculate the number of FOVs, that would have been recorded in the low-biomass samples, if they would have contained cells with good signals. The cell number counted within the few recorded FOVs was then considered to be counted within the corrected number of FOVs.

#### *Loss correction for air cell abundances*

These cell numbers were then corrected for losses due to re-aerosolization. Cell abundances reported within this manuscript are not corrected for reaerosolization and hence present a lower limit. The corrected values can be found in Dataset S1 and are summarised in Table 7.

Microbial cell abundances were corrected for losses due to re-aerosolization as described in (41, Equation **??**).

$$c_{air} = \frac{0.999998 \cdot N_0 \cdot e^{-0.7701t} - N(t)}{1.29853 \cdot e^{-0.7701t} - 1.29853} \cdot \frac{V_{col}}{v_{col}}$$

Where $c_{air}$: cell abundance in the sampled air [cells $m^{-3}$], $N_0$: cell abundance in the field blanks [cells $mL^{-1}$], $N(t)$: cell abundance in the sample [$mL^{-1}$], $t$: collection time [h], $v_{col}$: collection speed [$m^3$ $h^{-1}$], $V_{col}$: collection volume in Coriolis cone [mL]. $N_0$, $N(t)$ and $V_{col}$ refer to concentrations and volumes of the collection liquid, not the collected medium (air).

### Supporting Datasets

**ds1-samplelist_v2.xlsx –** Sample list including all relevant parameters discussed and visualized in this manuscript.
**ds2-16S-air-phylum-RSA.csv –** Phylum level relative sequence abundances of air samples. Last two columns show average and standard deviation over all samples.
**ds3-16S-water-phylum-RSA.csv –** Phylum level relative sequence abundances of water samples. Last two columns show average and standard deviation over all samples.
**ds4_test.xlsx –** Results of paired tests of water and air communities for selected clades of interest. First column: Clade name, Second column: Used test, Third column: V-value for paired Wilcoxon signed rank test, t-value for paired t-test, Rho for two-sided Spearman rank correlation, third column: p-value.
**ds5_biome_association.xlsx –** Assigned grouping of each lineage for each phylogenetic level as described in Tbl. 6. Microbial lineages were assigned to one of the respective grouping based on the relative sequence abundance difference in the two biomes (air and water) together with their prevalence, total relative sequence abundance and correlation between the RSA of the two biomes.
**ds6_ASVs.csv –** ASV by sample and taxonomy table as generated with the DADA2 workflow. Potential contaminant reads and ASVs were removed. This data set is the one presented within this manuscript, all ASVs in blank samples are zero after contamination removal, hence, this table does not contain the blank samples. Taxonomy was assigned using the SILVA database138.1 as described in the SI Appendix, Methods section.
**ds7_extractiontest.xlsx –** Overview of the results from the four tested DNA extraction protocols.
**ds8_testprime.xlsx –** 16S rRNA sequencing primer overview with their coverage of Bacteria, Archaea and Eukaryota depending on the number of mismatches as determined using TestPrime 1.0.
**ds9_dadaParams.xlsx –** Tested truncation and filtering parameters for DADA2 workflow and the number of reads passing through each step. Numbers are based on all samples of the sequencing run which includes samples not part of this study.
**ds10_contamremoval.xlsx –** Tested decontamination strategies. Rows 3 to 48 are single step approaches, rows 50 to 90 are combined approaches. Combination 1 and 2 are similar to the decontamination strategy used in (37), combination 3-5 use the low biomass function of the decontam package and differ in the way of subtracting/removing additional ASVs in the following steps. The percentage of ASVs/reads that are kept after each step are listed in the row of each step, the following row lists the percentage of ASVs/reads left of the original unfiltered data set.
**ds11_contaminants.xlsx –** List of potential contaminant clades used to filter sequencing data. First column indicates whether the respective clade was found to be a reagent or human contaminant.
**ds12_microscopy.xlsx –** Representative field of view from automated image recording for each sample. One field of view (image) has a width x height of 1388 x 1040 pixels. One pixel has a width/height of 0.1016 µm, hence, one image has a width x height of 141.0208 x 105.664 µm.
**ds13_contam-correlation.xlsx –** Correlations between the relative sequence abundances of clades considered potential reagent contaminants and the sample set.
**ds14_singlesampleBTs.pdf –** Three-day backward trajectories for each sample. Trajectories are shown for the air samples, unless a water sample was collected without a corresponding air sample, then the one backward trajectory belonging to the respective water sample is shown. Trajectories were truncated in case moderate rain occurred (more than 0.5 mm $h^{-1}$) as described in the Methods section.
**ds15-correlationsoverview.xlsx –** Correlations between cell abundances, DNA concentration, Richness and Inverse Simpson Diversity with latitude and surface ocean temperature for atmospheric and surface ocean samples. Pearsons Rho and p-values are given, for both, the whole dataset of air and water samples, as well as for subsets of the dataset based on the sampled longhurst provinces.
**ds16-TransmissionEfficiency-cpc-smps.csv –** Tubing configuration and transmission efficiency calculation results for CPC and SMPS, as calculated using the desktop version and Igor version of the particle loss calculator (42).
**ds17-smpsstats.xlsx –** Median, mean, first quartile, and third quartile of size bins from SMPS for the three air mass categories; marine, mixed, and terrestrial background, as shown in Fig. S9E.

### References


1. Abell, G. C. J. & Bowman, J. P. Ecological and biogeographic relationships of class Flavobacteria in the Southern Ocean. *FEMS microbiology ecology* **51**, 265–277, DOI: 10.1016/j.femsec.2004.09.001 (2005).

2. Meinhard, S., Glöckner, F.-O. & Amann, R. Different community structure and temperature optima of heterotrophic picoplankton in various regions of the Southern Ocean. *Aquatic Microb. Ecol.* **18**, 275–284 (1999).

3. Karner, M. B., DeLong, E. F. & Karl, D. M. Archaeal dominance in the mesopelagic zone of the Pacific Ocean. *Nature* **409**, 507–510, DOI: 10.1038/35054051 (2001).

4. Amano-Sato, C., Akiyama, S., Uchida, M., Shimada, K. & Utsumi, M. Archaeal distribution and abundance in water masses of the Arctic Ocean, Pacific sector. *Aquatic Microb. Ecol.* **69**, 101–112, DOI: 10.3354/ame01624 (2013).

5. Moss, J. A., Henriksson, N. L., Pakulski, J. D., Snyder, R. A. & Jeffrey, W. H. Oceanic Microplankton Do Not Adhere to the Latitudinal Diversity Gradient. *Microb. Ecol.* **79**, 511–515, DOI: 10.1007/s00248-019-01413-8 (2020).

6. Longhurst, A. R. Chapter 1 - toward an ecological geography of the sea. In *Ecological Geography of the Sea (Second Edition)*, DOI: https://doi.org/10.1016/B978-012455521-1/50002-4 (Academic Press, Burlington, 2007), second edition edn.

7. Zhou, J., Bruns, M. A. & Tiedje, J. M. DNA recovery from soils of diverse composition. *Appl. Environ. Microbiol.* **62**, 316–322, DOI: D-NLM:PMC167800EDAT-1996/02/01MHDA-1996/02/0100:01CRDT-1996/02/0100:00PST-ppublish (1996). arXiv:1408.1149.

8. R Core Team. *R: A Language and Environment for Statistical Computing*. R Foundation for Statistical Computing (2023).

9. Dowle, M. & Srinivasan, A. *data.table: Extension of ‘data.frame‘* (2023). R package version 1.14.8.

10. Wickham, H., François, R., Henry, L., Müller, K. & Vaughan, D. *dplyr: A Grammar of Data Manipulation* (2023). R package version 1.1.2.

11. Hamilton, N. E. & Ferry, M. ggtern: Ternary diagrams using ggplot2. *J. Stat. Software, Code Snippets* **87**, 1–17, DOI: 10.18637/jss.v087.c03 (2018).

12. Wickham, H. *ggplot2: Elegant Graphics for Data Analysis* (Springer-Verlag New York, 2016).

13. Neuwirth, E. *RColorBrewer: ColorBrewer Palettes* (2022). R package version 1.1-3.

14. Oksanen, J. *et al.* *vegan: Community Ecology Package* (2022). R package version 2.6-4.

15. Callahan, B. J. *et al.* Dada2: High-resolution sample inference from illumina amplicon data. *Nat. Methods* **13**, 581–583, DOI: 10.1038/nmeth.3869 (2016).

16. Larsson, J. & Gustafsson, P. A case study in fitting area-proportional Euler diagrams with ellipses using eulerr. In *Proceedings of International Workshop on Set Visualization and Reasoning*, vol. 2116, 84–91 (CEUR Workshop Proceedings, Edinburgh, United Kingdom, 2018).

17. Wickham, H. *et al.* Welcome to the tidyverse. *J. Open Source Softw.* **4**, 1686, DOI: 10.21105/joss.01686 (2019).

18. Wickham, H. Reshaping data with the reshape package. *J. Stat. Softw.* **21**, 1–20 (2007).

19. Constantin, A.-E. & Patil, I. ggsignif: R package for displaying significance brackets for ’ggplot2’. *PsyArxiv* DOI: 10.31234/osf.io/7awm6 (2021).

20. Millard, S. P. *EnvStats: An R Package for Environmental Statistics* (Springer, New York, 2013).

21. Kassambara, A. *ggpubr: ’ggplot2’ Based Publication Ready Plots* (2023). R package version 0.6.0.

22. Paradis, E. & Schliep, K. ape 5.0: an environment for modern phylogenetics and evolutionary analyses in R. *Bioinformatics* **35**, 526–528, DOI: 10.1093/bioinformatics/bty633 (2019).

23. Gehlenborg, N. *UpSetR: A More Scalable Alternative to Venn and Euler Diagrams for Visualizing Intersecting Sets* (2019). R package version 1.4.0.

**24.** Dusa, A. *venn: Draw Venn Diagrams* (2022). R package version 1.11.

**25.** De Cáceres, M. & Legendre, P. Associations between species and groups of sites: indices and statistical inference. *Ecology* **90**, 3566–3574, DOI: 10.1890/08-1823.1 (2009).

**26.** Schauberger, P. & Walker, A. *openxlsx: Read, Write and Edit xlsx Files* (2023). R package version 4.2.5.2.

**27.** Ushey, K., Allaire, J., Wickham, H. & Ritchie, G. *rstudioapi: Safely Access the RStudio API* (2022). R package version 0.14.

**28.** van den Boogaart, K. G., Tolosana-Delgado, R. & Bren, M. *compositions: Compositional Data Analysis* (2023). R package version 2.0-6.

**29.** Murrell, P. & Wen, Z. *gridGraphics: Redraw Base Graphics Using 'grid' Graphics* (2020). R package version 0.5-1.

**30.** Davis, N. M., Proctor, D., Holmes, S. P., Relman, D. A. & Callahan, B. J. Simple statistical identification and removal of contaminant sequences in marker-gene and metagenomics data. *Microbiome* **6**, 1–14 (2018).

**31.** Stauffer, R., Mayr, G. J., Dabernig, M. & Zeileis, A. Somewhere over the rainbow: How to make effective use of colors in meteorological visualizations. *Bull. Am. Meteorol. Soc.* **96**, 203–216, DOI: 10.1175/BAMS-D-13-00155.1 (2009).

**32.** Massicotte, P. & South, A. *rnaturalearth: World Map Data from Natural Earth* (2023). R package version 0.3.3.

**33.** Mallick, H., Rahnavard, A. & McIver, L. J. *MaAsLin 2: Multivariable Association in Population-scale Meta-omics Studies.* (2020). R/Bioconductor package.

**34.** Wickham, H., Vaughan, D. & Girlich, M. *tidyr: Tidy Messy Data* (2023). R package version 1.3.0.

**35.** Quast, C. *et al.* The SILVA ribosomal RNA gene database project: improved data processing and web-based tools. *Nucleic Acids Res.* **41**, D590–D596, DOI: 10.1093/nar/gks1219 (2012).

**36.** Yilmaz, P. *et al.* The SILVA and "All-species Living Tree Project (LTP)" taxonomic frameworks. *Nucleic Acids Res.* **42**, D643–D648, DOI: 10.1093/nar/gkt1209 (2014).

**37.** Lang-Yona, N. *et al.* Terrestrial and marine influence on atmospheric bacterial diversity over the north Atlantic and Pacific Oceans. *Commun. Earth & Environ.* **3**, 121, DOI: 10.1038/s43247-022-00441-6 (2022).

**38.** Salter, S. J. *et al.* Reagent and laboratory contamination can critically impact sequence-based microbiome analyses. *BMC Biol.* **12**, 1–12, DOI: 10.1186/s12915-014-0087-z (2014).

**39.** Barton, H., Taylor, N., Lubbers, B. & Pemberton, A. DNA extraction from low-biomass carbonate rock: An improved method with reduced contamination and the low-biomass contaminant database. *J. Microbiol. Methods* **66**, 21–31, DOI: 10.1016/j.mimet.2005.10.005 (2006).

**40.** Brüwer, J. D. *et al.* In situ cell division and mortality rates of SAR11, SAR86, *Bacteroidetes*, and *Aurantivirga* during phytoplankton blooms reveal differences in population controls. *mSystems* **e01287-22**, DOI: 10.1128/msystems.01287-22 (2023).

**41.** Mayol, E., Jiménez, M. A., Herndl, G. J., Duarte, C. M. & Arrieta, J. M. Resolving the abundance and air-sea fluxes of airborne microorganisms in the North Atlantic Ocean. *Front. Microbiol.* **5**, 557, DOI: 10.3389/fmicb.2014.00557 (2014).

**42.** von der Weiden, S.-L., Drewnick, F. & Borrmann, S. Particle Loss Calculator – a new software tool for the assessment of the performance of aerosol inlet systems. *Atmospheric Meas. Tech.* **2**, 479–494, DOI: 10.5194/amt-2-479-2009 (2009).

**43.** Legendre, P. & Gallagher, E. D. Ecologically meaningful transformations for ordination of species data. *Oecologia* **129**, 271–280, DOI: 10.1007/s004420100716 (2001).

**44.** Anderson, M. J., Ellingsen, K. E. & McArdle, B. H. Multivariate dispersion as a measure of beta diversity. *Ecol. Lett.* **9**, 683–693, DOI: 10.1111/j.1461-0248.2006.00926.x (2006).

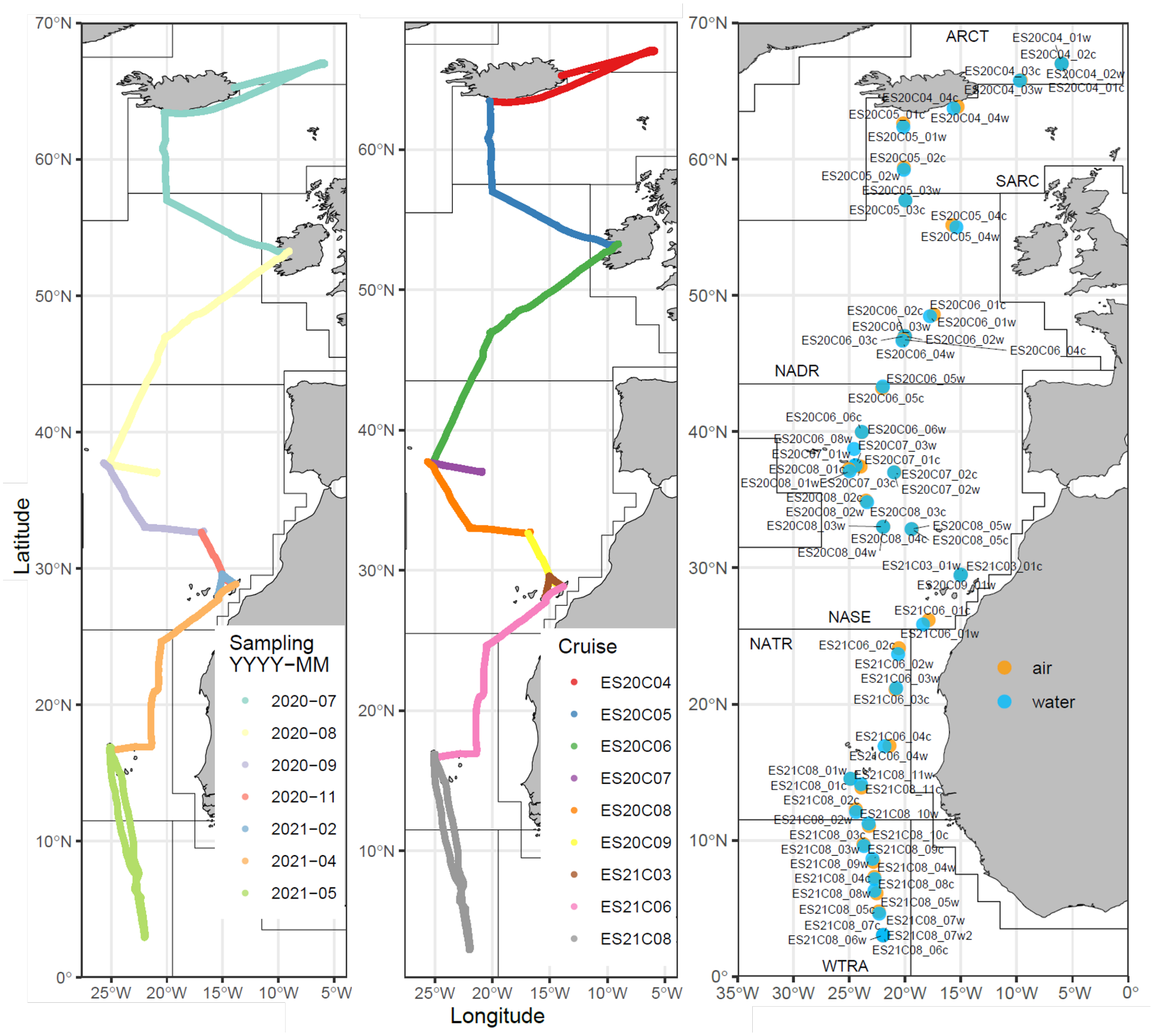


**Figure 1.** Left: Cruise track colored by cruise year and month (YYYY-MM). Middle: Cruise track colored by unique cruise identifier. Right: Air and water sampling locations along the latitudinal transect.

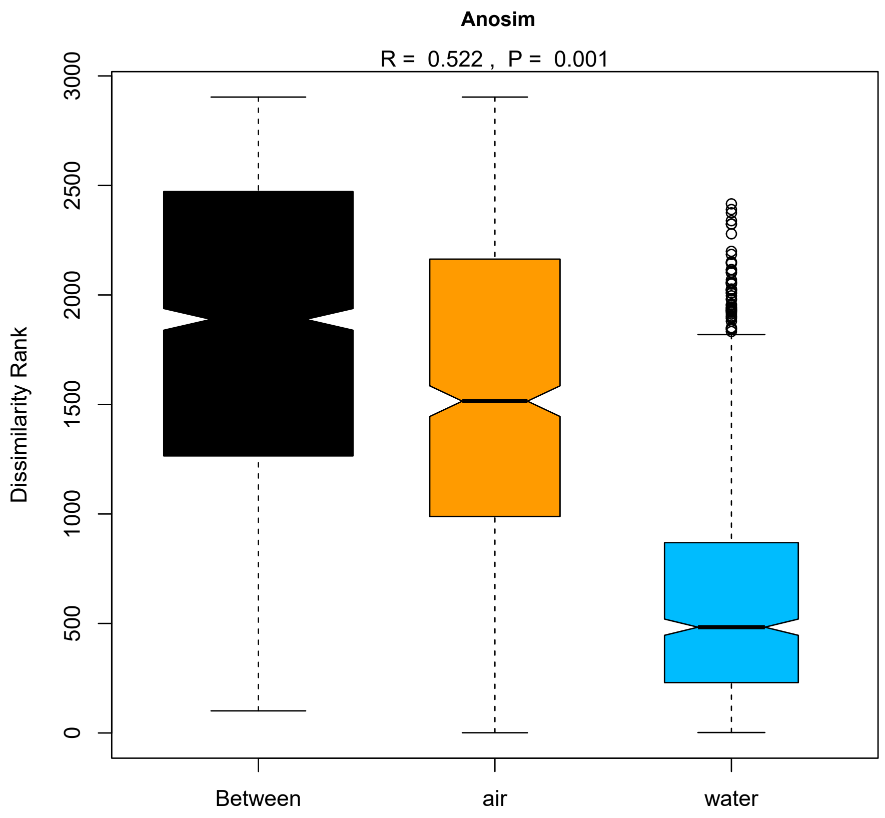


**Figure 2.** Anosim Dissimilarity Ranks between air and water samples and within groups.

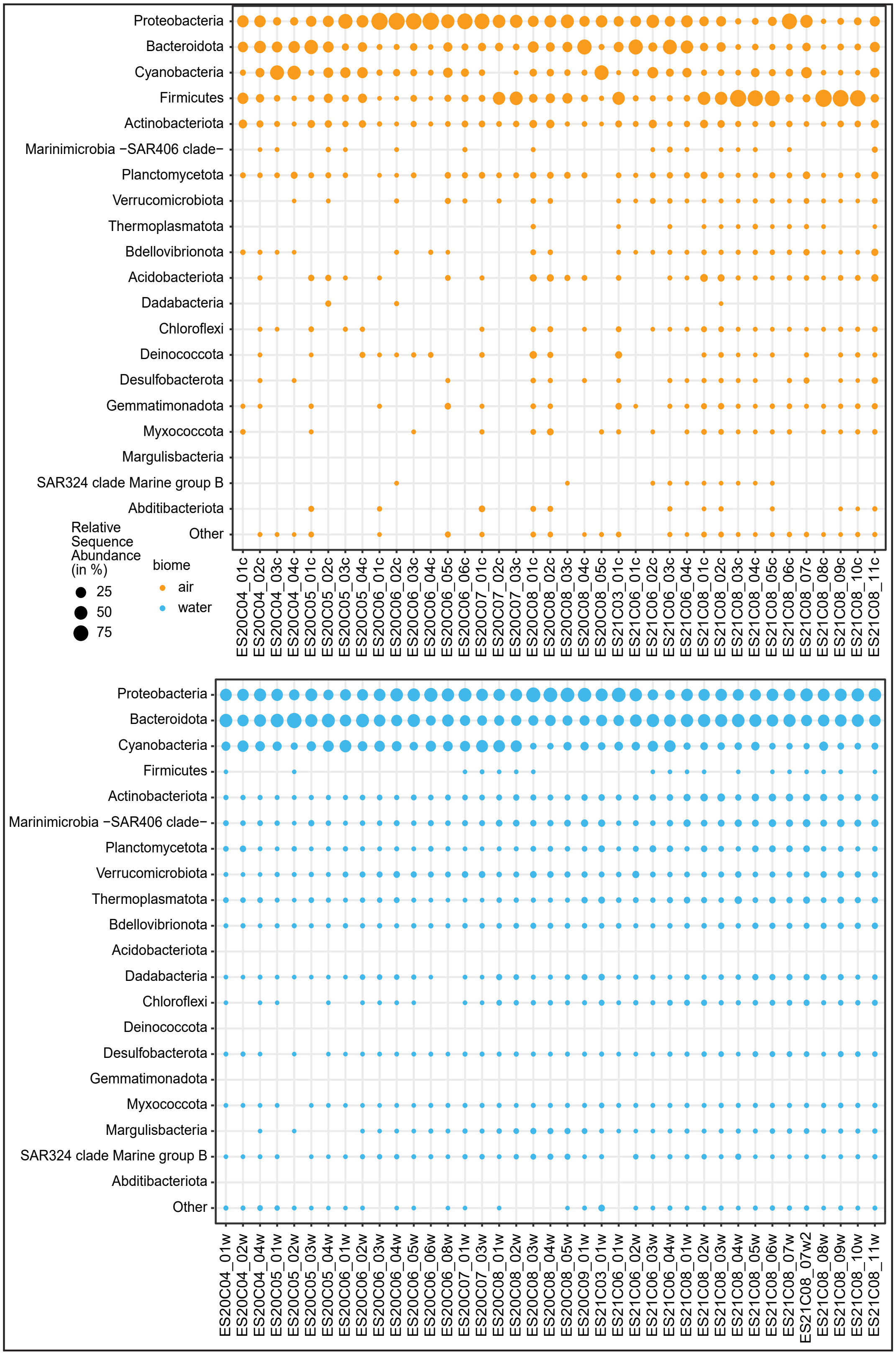


**Figure 3.** Phylum level clades air (orange) and marine (blue) samples. Bubble size is scaled with relative sequence abundance of each phylum. Phyla are ordered by decreasing average of their relative sequence abundance in the whole data set. Samples within groups are ordered chronologically.

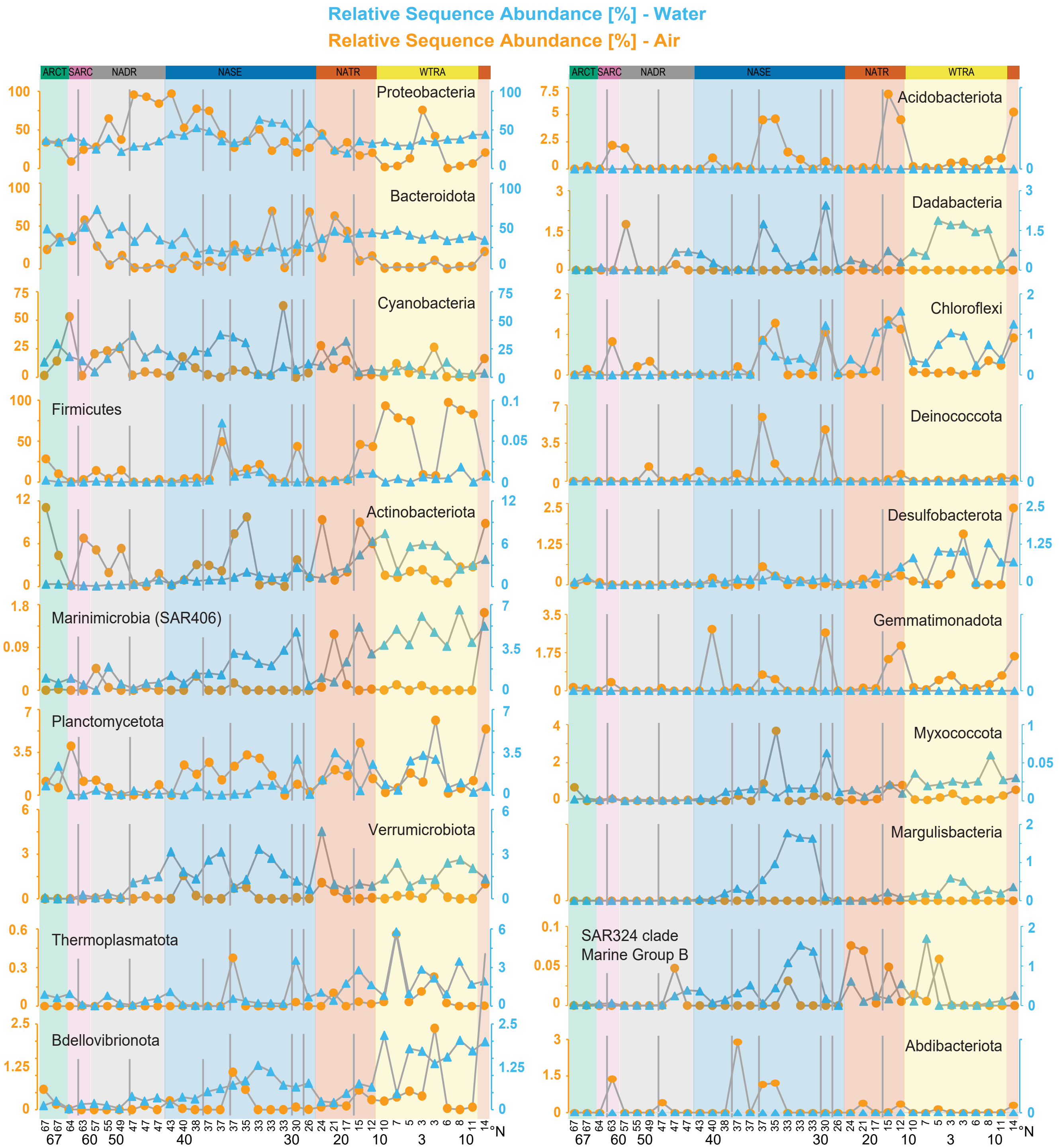


**Figure 4.** Relative sequence abundance of 20 most abundant phyla found in air (orange) and water (blue) in %. If all samples in the biome had a RSA of zero, the corresponding y-axis only displays a zero. Phylum with highest relative sequence abundance (Proteobacteria) top left, Phylum with rank 11 (Acidobacteriota) on the top right, RSAs decreasing within one column. Within one plot, samples are ordered chronologically from left to right, following the latitudinal transect. Vertical lines indicate breaks between cruises. Samples within one cruise are usually one day apart. Longhurst provinces are highlighted (green: Atlantic Arctic (ARCT), purple: Atlantic sub-Arctic (SARC), grey: North Atlantic Drift (NADR), blue: North Atlantic subtropical gyre (NASE), orange: North Atlantic Tropical Gyral (NATR), yellow: Western Tropical Atlantic (WTRA)).

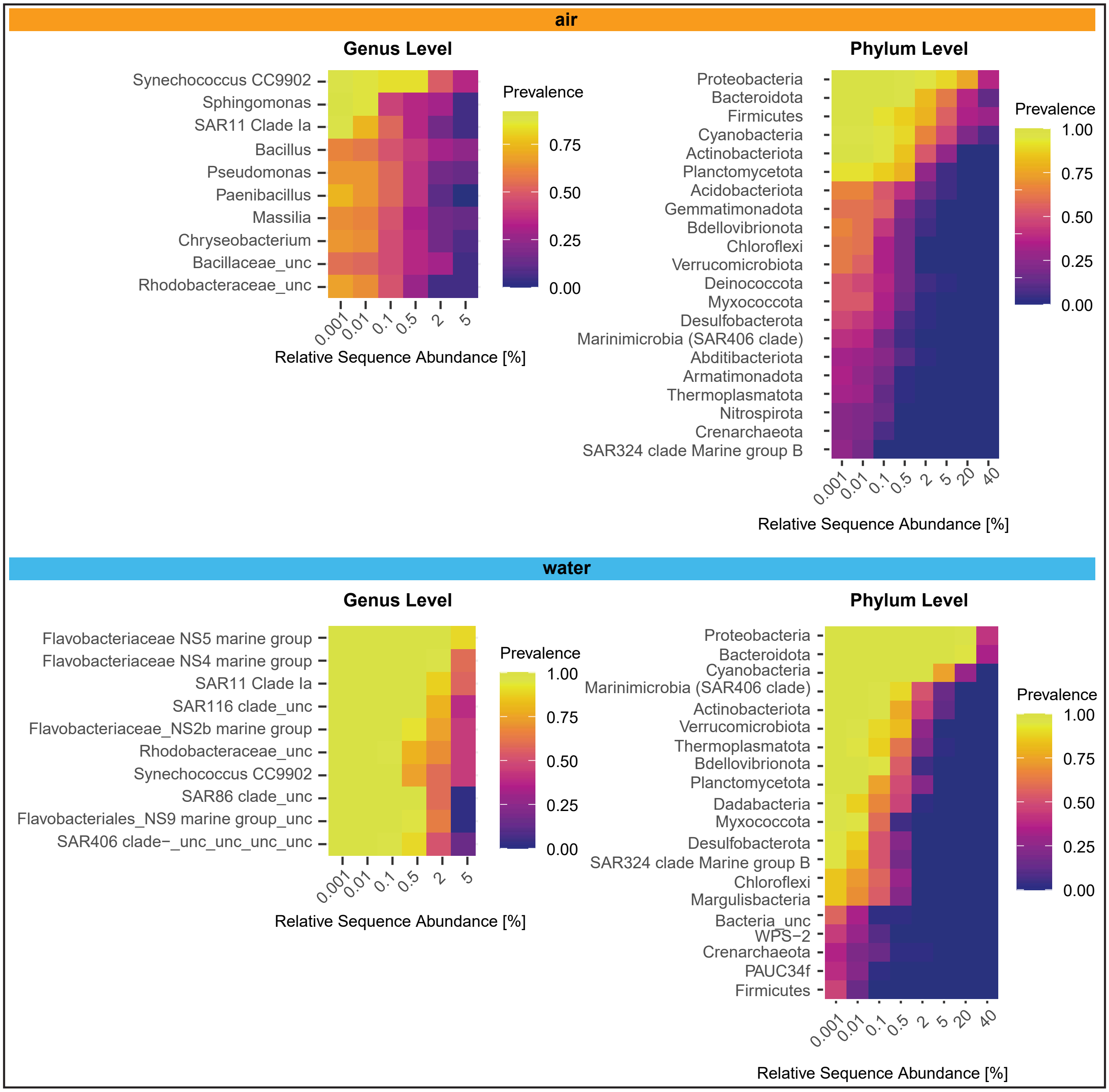


**Figure 5.** Top members of a potential core microbiome of the open ocean air and surface water community based on their relative abundance and prevalence in samples from the North East Atlantic Ocean between 67° N and 3° N.

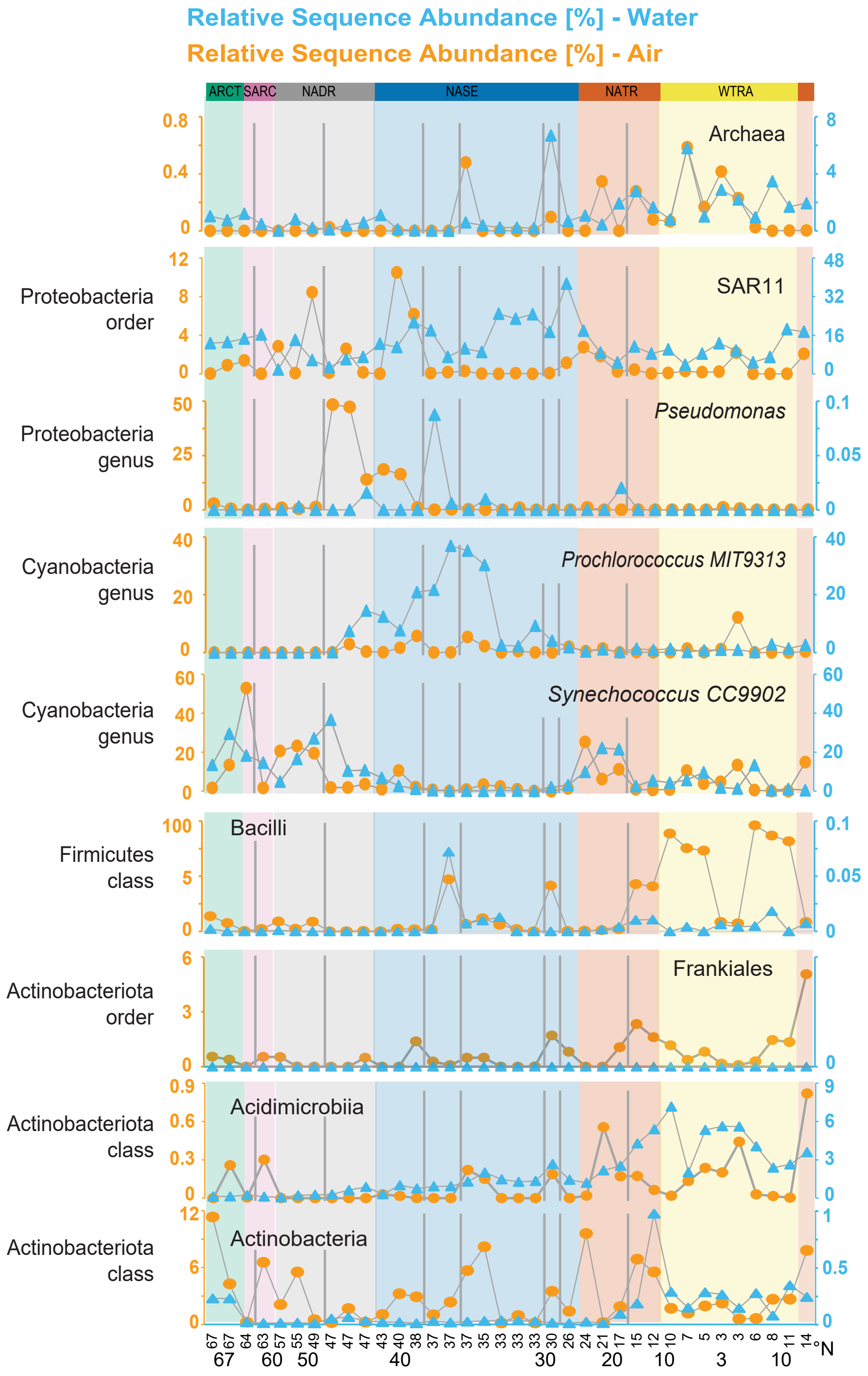


**Figure 6.** Relative sequence abundance (RSA) of microbial lineages discussed in the main text for air (orange) and water (blue) in %. If all samples in the biome had a RSA of zero, the corresponding y-axis only displays a zero. Samples are ordered chronologically from left to right, following the latitudinal transect. Vertical lines indicate breaks between cruises. Samples within one cruise are usually one day apart. Longhurst provinces are highlighted (Green: Atlantic Arctic (ARCT), purple: Atlantic sub-Arctic (SARC), grey: North Atlantic Drift (NADR), blue: North Atlantic subtropical gyre (NASE), orange: North Atlantic Tropical Gyral (NATR), yellow: Western Tropical Atlantic (WTRA)).

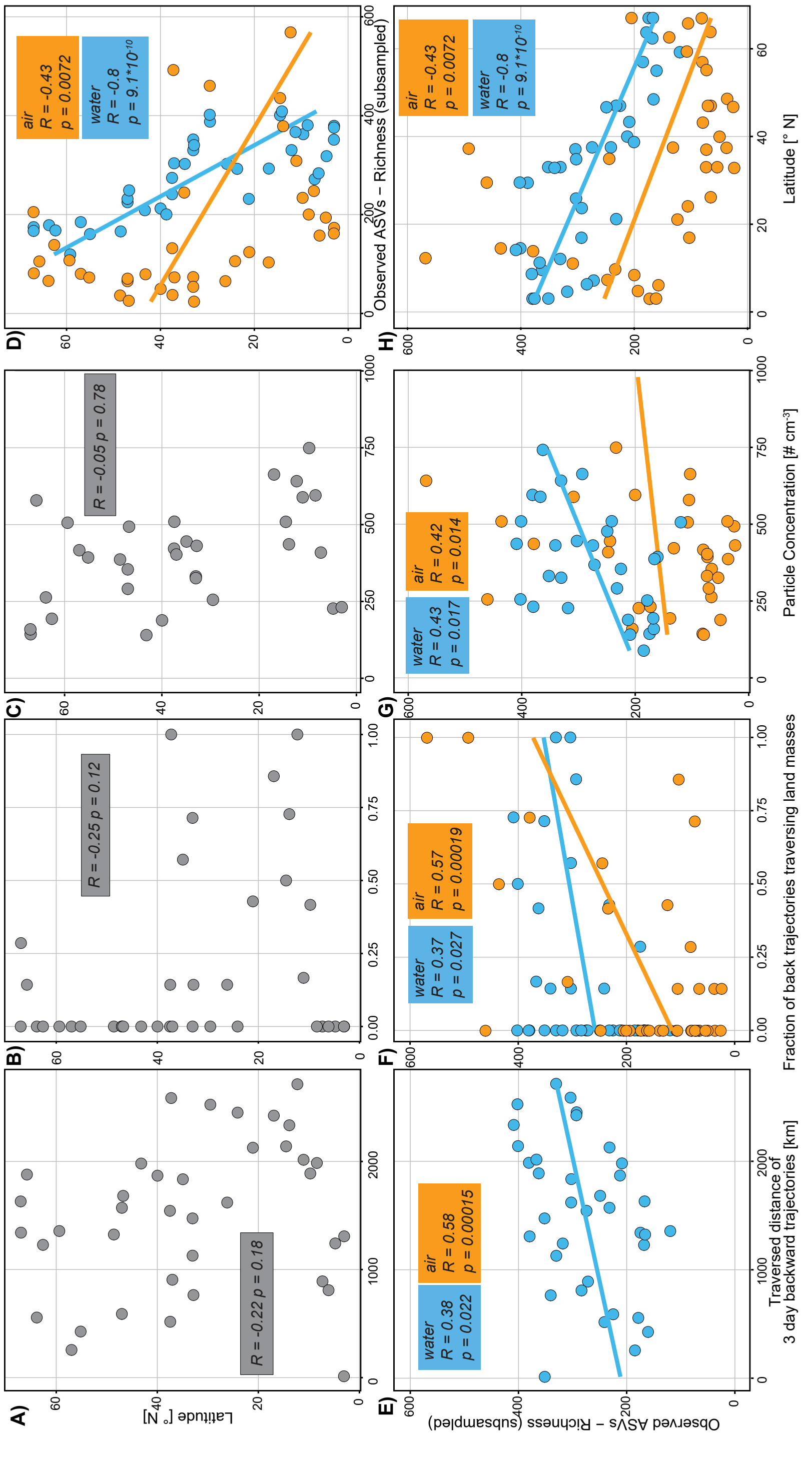


**Figure 7.** Dependence of air (blue) and water (orange) community richness on air mass origin, particle concentration and latitude. A) The traversed distance of the calculated 3-day backward trajectories does not correlate with latitude. B) The fraction of 3-day backward trajectories traversing land masses before arrival at the sampling location does not correlate with latitude. C) The particle concentration observed at the sampling site does not correlate with latitude. D) The observed community richness of air and water negatively correlates with the sampled latitude [° N]. E-G) The traversed distance by the calculated 3-day backward trajectories, the fraction of trajectories traversing land masses and the measured particle concentration at site, positively correlate with the observed community richness in air and water. R: Pearson's *R* together with the significance ($p$) as calculated with ggpubr::stat_cor(). Lines show linear regression result as calculated using geom_smooth(method='lm',formula=y~x). C & G one outlier with particle concentration >1000 particles per $cm^{-3}$ not shown but included in the calculation of the correlation and regression.

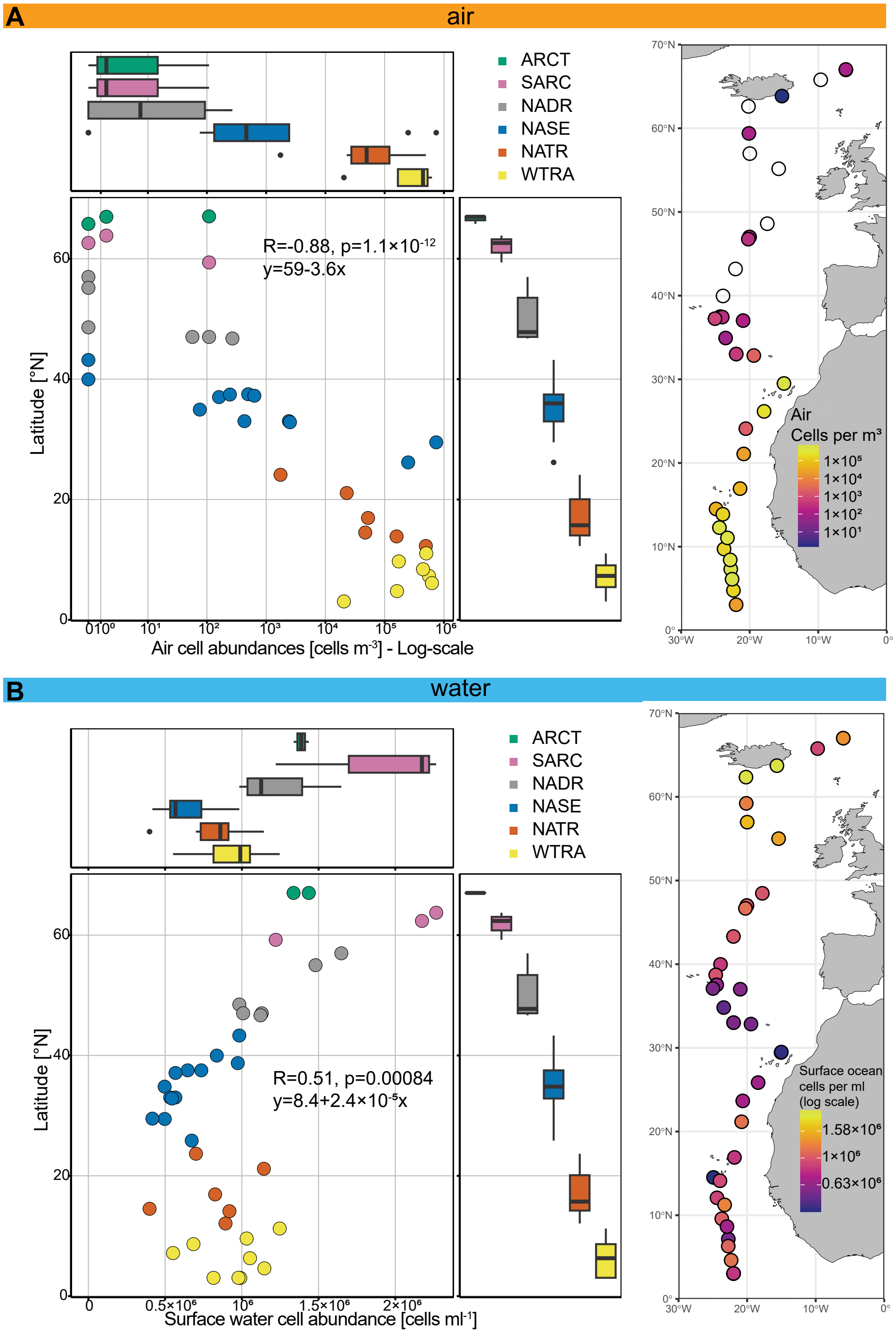


**Figure 8.** Latitude [° N] vs cell abundance in the North East Atlantic. Samples are colored-coded by their association with the 6 sampled oceanic provinces (Table 2). A: Left: Latitude vs air cell abundance on a logarithmic scale. Cell abundances show a significant negative correlation with latitude. Right: Map with atmosphere samples for cell enumeration colored by the resulting cells per cubic meter. Logarithmic scale. Zero values are empty circles. B: Left: Latitude vs surface ocean cell abundances. Cell abundances show an overall positive correlation with latitude, with a minimum at around 30° N. Right: Map with surface ocean samples taken for cell enumeration colored by the resulting cells per mL. Logarithmic scale.

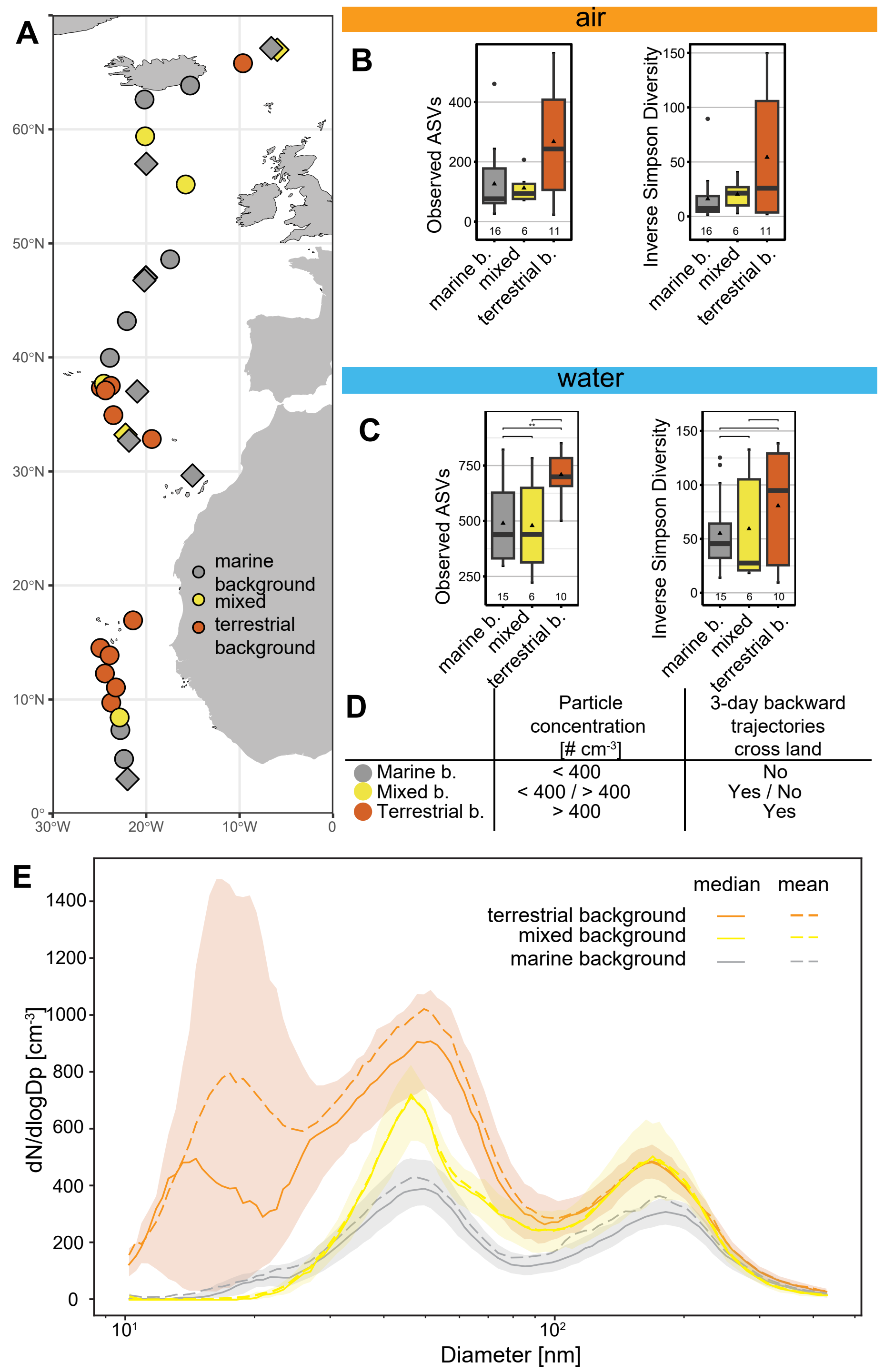


**Figure 9.** Samples grouped by potential air mass origin. A: Samples along the latitudinal transect and their assignment to the respective group. Circles are samples taken while cruising, diamonds are samples taken at stations. Where needed, circles have been stacked to display all background conditions at their approximate location. B&C: Boxplots of alpha diversity of the air (B) and surface ocean water (C). ** = p<0.01. D: Summarized grouping definitions as defined in the main manuscript. E: Particle number size distributions (PNSDs)for the three air mass origin groups, normalized by bin width (dN/dlogDp), showing an increase in small particles during sampling periods classified as terrestrial background, in contrast to mixed and marine background conditions. Straight lines show median PNSDs, dashed lines show mean PNSDs, shaded areas shows the range between the first and third quartiles. Note the logarithmic x axis scale.

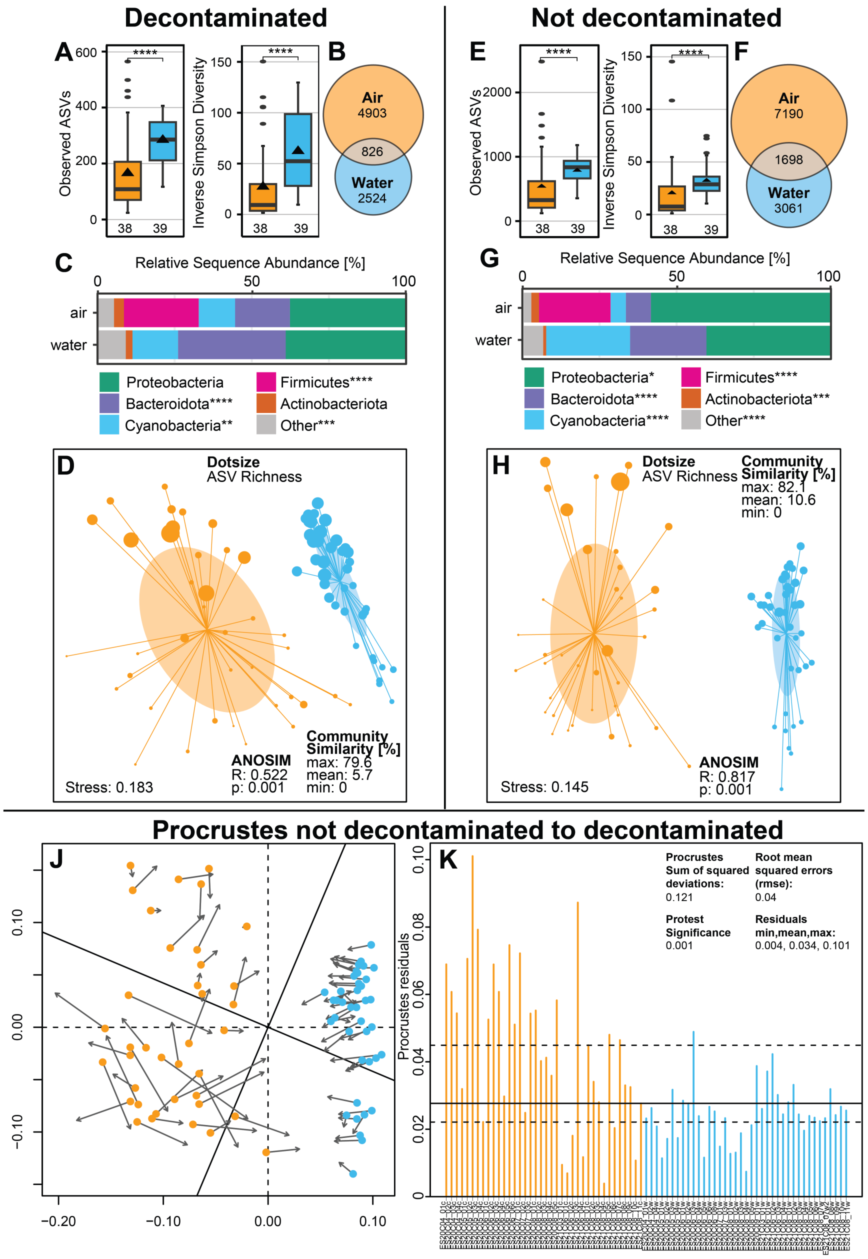


**Figure 10.** Microbial diversity of lower atmosphere and surface ocean communities of the North East Atlantic Ocean of the decontaminated data set (A-D), and the dataset without removal of any microbial clades (except mitochondrial and chloroplast DNA, E-H). A,E: Microbial alpha diversity of air (orange, n= 38) and water (blue, n = 39) samples. Boxplots show mean (triangle), median (middle line), upper and lower quartile (top and lower line). Whiskers show maximum data points within upper/lower quartile plus 1.5 times the interquartile range (IQR). Dots represent outlier values that exceed 1.5×IQR. Groups were tested using a Wilcoxon rank sum test, ****: p<0.0001. B,F: Scaled Venn Diagram showing unique and shared ASVs between surface ocean and air, corrected for group size. C,G: Relative sequence abundances of the five most abundant phyla. Differences between air and water were tested using a Wilcoxon rank sum test, ****: p<0.0001, ***: p<0.001, **: p<0.01, *: p<0.05. D,H: Non-metric multidimensional scaling (NMDS) plot of air (orange) and water (blue) communities. The closer two dots are, the more similar are their underlying communities. Dot size scales with sample richness, oval shapes depict one standard deviation of the centroid (weighted average within-group-distance). Lines connect each dot to the centroid of the tested group. J: Procrustes (least-squares orthogonal mapping) of NMDS ordinations for not-decontaminated and decontaminated dataset. K: Residuals showing the deviations between two data points of the procrustes fit. Horizontal lines are 25, 50, and 75 % quantiles of the residuals. Protest procrustean randomization test significance given using vegan::protest() with permutations set to 1000, note that significance in this case can not be less than 0.001.

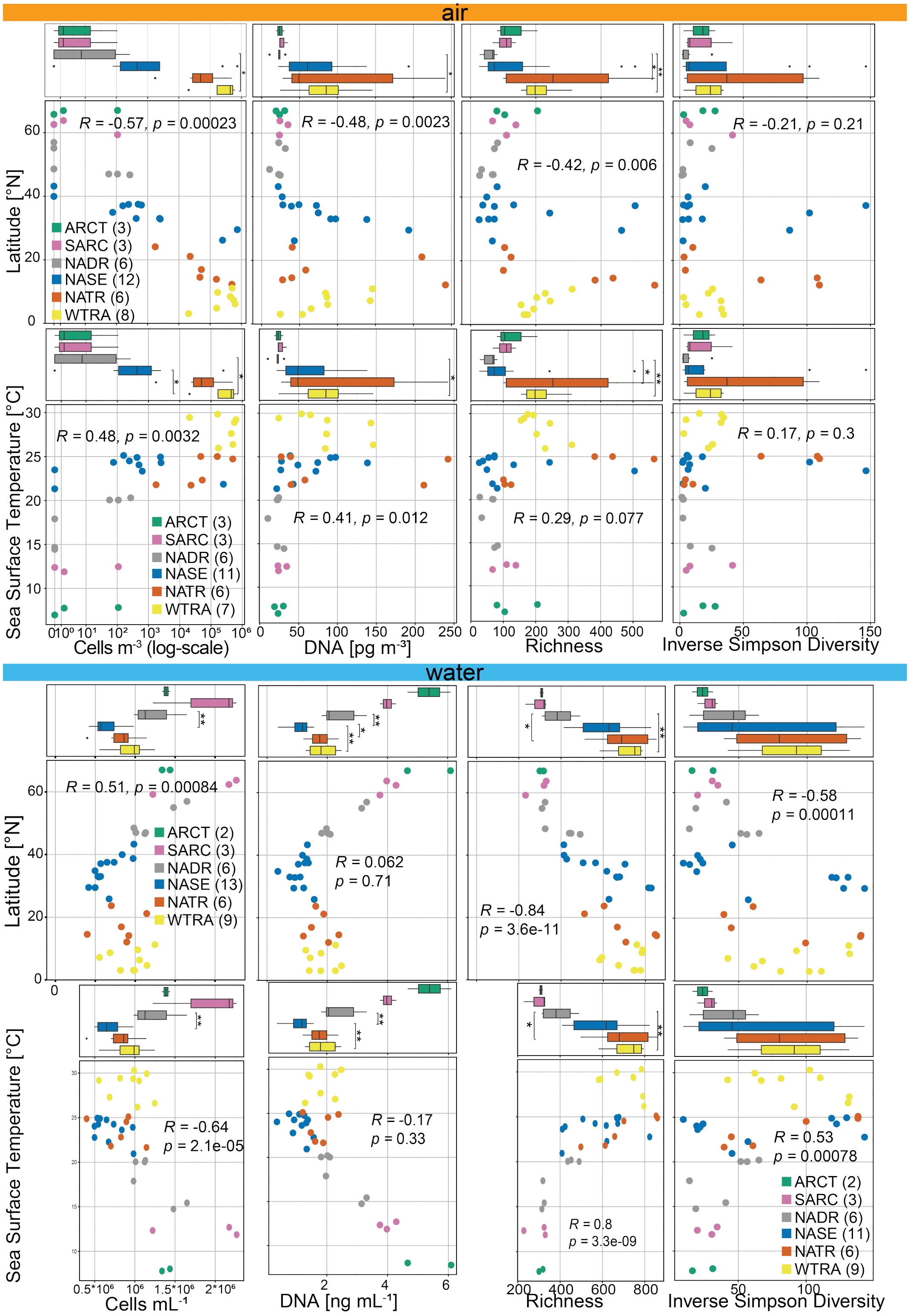


**Figure 11.** Changes in cell abundances, DNA concentration, Richness and Inverse Simpson Diversity of air and surface water communities along Latitude and Sea Surface Temperature, grouped by oceanic provinces (6). Differences between longhurst provinces were tested using a Wilcoxon rank sum test with bonferroni correction, **: p<0.01, *: p<0.05. Pearson correlation Rho and p are given. Correlations for each subgroup are summarized in SI Dataset S15. Group size is given for each row. Green: Atlantic Arctic (ARCT), purple: Atlantic sub-Arctic (SARC), grey: North Atlantic Drift (NADR), blue: North Atlantic subtropical gyre (NASE), orange: North Atlantic Tropical Gyral (NATR), yellow: Western Tropical Atlantic (WTRA). DNA: 1 outlier in WTRA not shown (16.7 ng mL$^{-1}$ DNA, sample ES21C08_08w).

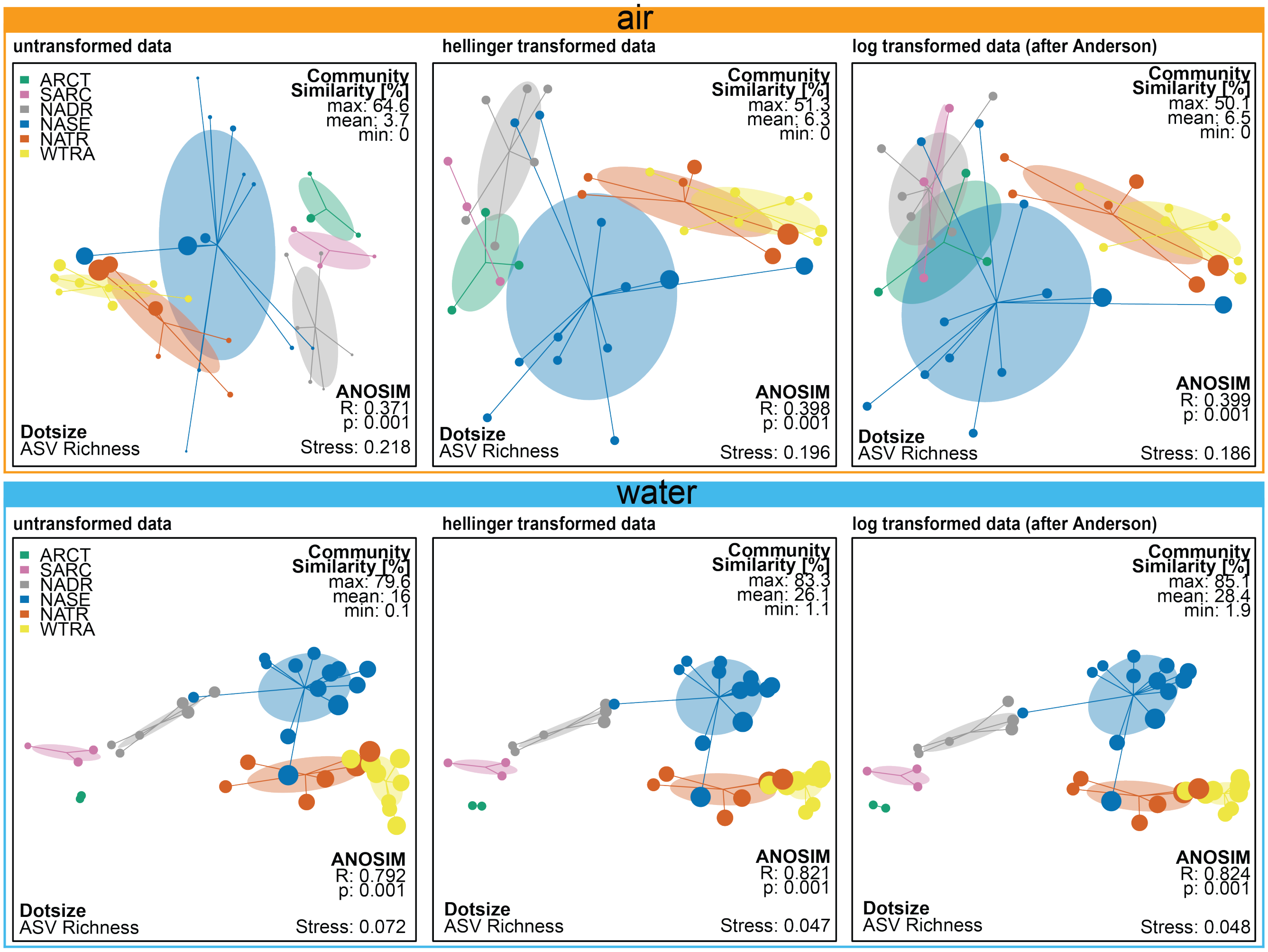


**Figure 12.** NMDS of air (top) and surface ocean (bottom) microbial communities, colored by oceanic provinces. Dots are colored by oceanic provinces and scaled by ASV richness. The data was either un-transformed before calculating the NMDS (left), or hellinger transformed (vegan::decostand(method ="hellinger"), middle, 43) or log transformed (vegan::decostand(method ="log"), right, 44) to investigate the effect of transformations on the stress values.

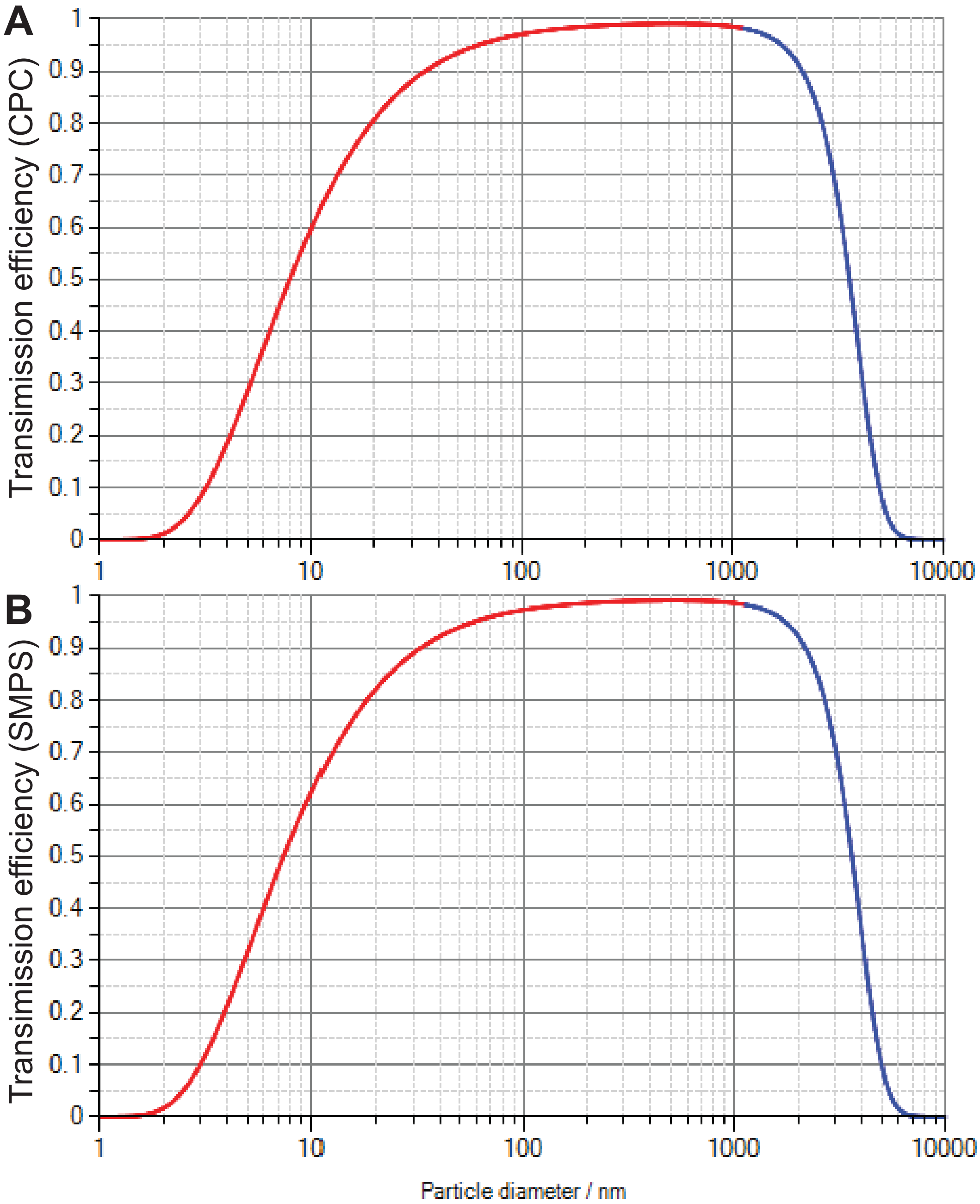


**Figure 13.** A) Transmission efficiency for particles between 1 nm and 10 µm particle size for the TSI CPC 3787. B) Transmission efficiency for particles between 1 nm and 10 µm particle size for the TSI SMPS. Transmission efficiency was calculated using the Particle loss calculator (42). Data outside formula validity is marked in red as calculations with an angle of enlargement larger than 4° result in an undererstimation of the occurring particle loss.

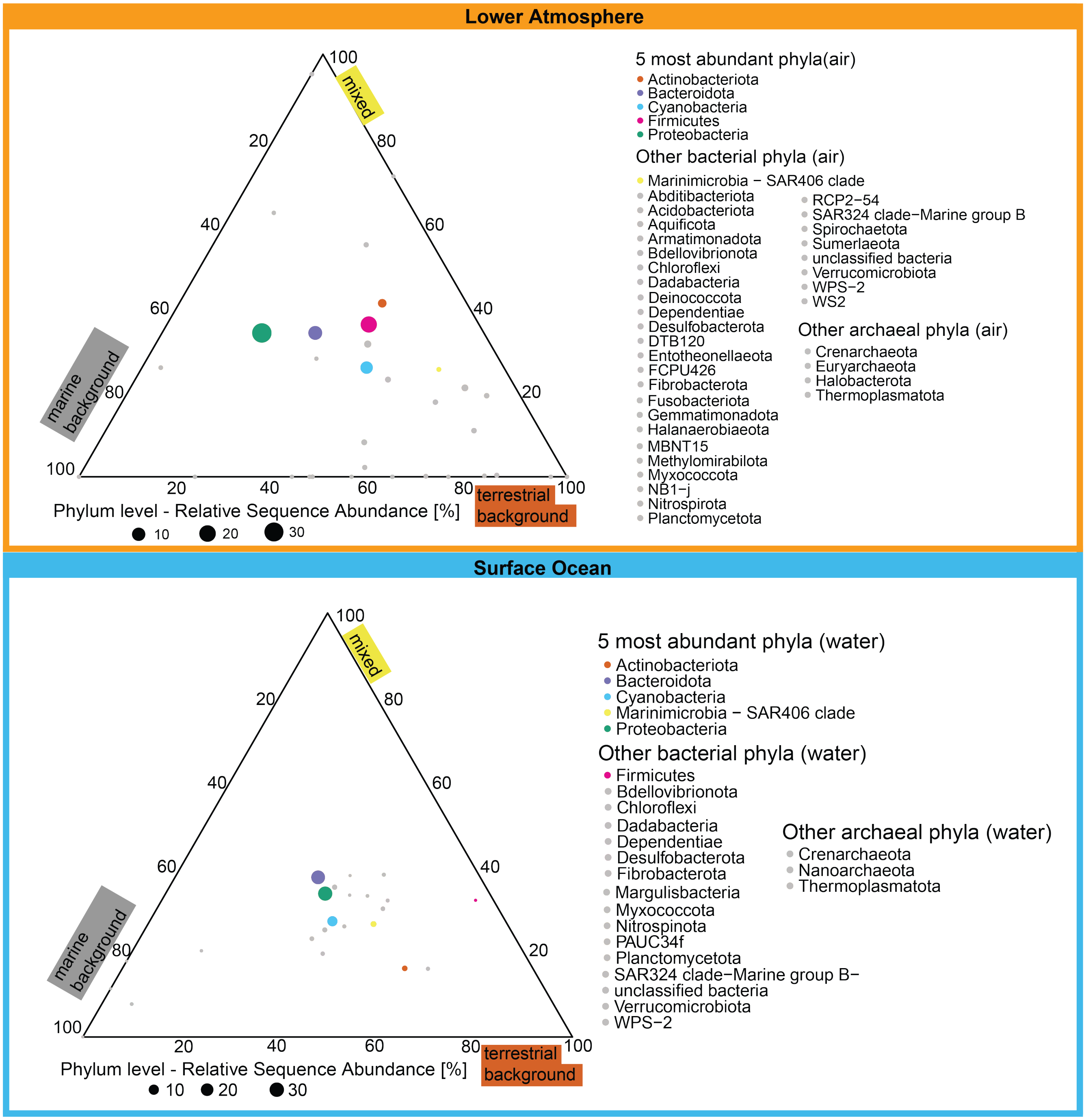


**Figure 14.** Ternary plots of air (top) and surface water (bottom) community composition under terrestrial, marine and mixed conditions. Closed circles represent taxonomy at phylum level, circle size represents relative sequence abundance of the respective taxon averaged across all samples of the respective biome. Circle position indicates the distribution across the conditions. Top 5 abundant phyla are distinct colors, while the remaining phyla ("Other") are grey.
